# Chronological Aging Stiffens Cells, Alters Mitochondria, and Impairs Mitochondrial Mechanical Responsiveness in Mesenchymal Stem Cells

**DOI:** 10.64898/2026.02.24.707807

**Authors:** Omar Faruq, Nina Nikitina, Scott Birks, Calvin L Jones, Matthew Goelzer, Sean Howard, Anamaria Zavala, Nilufar Ali, Gunes Uzer

## Abstract

Mesenchymal stem cells (MSCs) can differentiate into osteoblasts and adipocytes, play a critical role for maintaining bone homeostasis. Although, aging impairs MSC function and contributes to several complications, the effects of aging on subcellular structure and related gene expression need further investigation. Here we established an *in vitro* system to study MSCs isolated from bones of 5, 12, and 24 months old mice which showed significantly decreased bone volume and exercise performance with age. RNA sequencing revealed downregulation of genes related to cell-matrix interactions, cell metabolism, and division. Functionally MSCs extracted from 24mo showed significantly increased adipogenesis and reduced osteogenesis compared to 5mo, which was accompanied by reduced levels of cell proliferative marker Ki67 and increased expression of senescence protein p16. Data-driven segmentation of F-actin architecture revealed no cell-wide or nuclei-associated alteration, while 24mo MSCs showed decreased nuclear volume and increased spreading and higher nuclear stiffness compared to 5mo. Mitochondria from 12mo and 24mo showed increased length and volume of fibers when compared to 5mo, which was accompanied by gene expression changes associated with mitochondrial inflammation and oxidative phosphorylation. To test the functional consequences, MSCs were subjected to mechanical stress for 72 hours using 90Hz, 07g low-intensity vibration (LIV). While 5mo MSCs showed a robust fusing of individual mitochondrial fibers, LIV response of 12mo and 24mo MSCs were progressively less, indicating an already stressed mitochondria compromised to mechanical challenge. In summary, this study has established an in vitro assay system, revealing age-associated impairments in differentiation, mitochondrial function, and mechanotransduction capacity.

## 1 Introduction

Aging is an inevitable physiological process characterized by a progressive decline in the function of all organs and the onset of various dysfunctions contributing to approximately two-thirds of the 150,000 deaths that occur globally each day (Niccoli & Partridge, 2012). Notably aging is associated with decreased musculoskeletal competence, decreasing mobility and increasing the risk of injuries (Nguyen el al. 2025). A consequence of aging on the musculoskeletal system is decreased bone quality which leads to increased bone fragility and increased fatty infiltration of bone. Age-associated bone mass decline and increased fatty infiltration is in-part linked to the impaired function of mesenchymal stem cells (MSCs), which are capable of differentiating into osteoblasts and adipocytes (Pagnotti et al. 2019). The exhaustion of MSCs during aging is also a recognized hallmark of skeletal aging, leading to a reduced regenerative capacity in skeletal tissues (Zhou et al. 2008). The aging bone marrow microenvironment contributes to a decline in both the number and functional capacity of MSCs, including alterations in their differentiation potential (Bellantuono et al. 2009). Several studies have reported reduced proliferative capacity in cells from aged donors across multiple species (Stolzing et al. 2008). Additionally, research has shown that MSCs from older individuals exhibit increased cellular senescence compared to those from younger individuals (Wagner et al. 2008). Aging has also been associated with an increased adipogenic potential and a corresponding decrease in osteogenic differentiation (Moerman et al. 2004).

Despite characterization of these functional changes, how aging effects cellular structure is less understood. As mechanical forces play a critical role in regulating a number of biological processes, including cell migration, adhesion and differentiation (Leight et al. 2012), age-related changes in cellular form is critical for proper function of organs and tissue systems (Discher et al. 2009). Indeed, aging gradually impairs the mechanical properties of cells, affecting various aspects of mechanosignaling at both molecular and subcellular levels including changes in alterations in mitochondrial function, cytoskeletal architecture, and nuclear morphology (Han et al 2025, Kim et al. 2022). In this way, our study was aimed at analyzing the effects of aging on primary MSCs extracted from female C57BL/6 mice at both the subcellular and molecular levels, and to determine whether aged MSCs retain the ability to sense and respond to mechanical forces. MSCs were harvested from the bone marrow of mice aged 5, 12, and 24 months, and cultured *in vitro* to examine age-related changes in MSC characteristics. We performed RNA sequencing (RNA-seq), identified proliferative and differentiation characteristics. These assays were followed by structural analysis of F-actin cytoskeleton, nucleus, mitochondria and mechanical analysis of live cells. Finally, changes in mitochondrial structure in response to mechanical challenge was evaluated in response to Low Intensity Vibration (LIV), indicating an age-related decrease in cell’s ability to respond to mechanical stress.

## 2 Materials and Method

### 2.1 Animals

C57BL/6J female mice from three different age groups were obtained from NIH Aged Rodent Colonies (MSCs isolation) and Jackson laboratory, stock number 000664 (Micro-CT and wheel running) and housed in separately ventilated cages under controlled conditions. All animals were provided with appropriate access to food and water, and the study was conducted in compliance with institutional guidelines. All procedures were approved by the Boise State University Institutional Animal Care and Use Committee.

### 2.2 Micro-CT analysis

Proximal tibiae (n=10 to 12mice/group) were stored frozen in PBS-soaked gauze prior to scanning and were scanned immersed in PBS using an X-ray microtomograph (SkyScan 1172F). The scanning settings were as follows: power of 55 kV/181 μA, a voxel size of 10.33 µm³, and an integration time of 230 ms. The volume of interest for trabecular quantitative analysis began 10 slices distal to the proximal physis and extended 1 mm. The cortical volume of interest started 2.15 mm distal to the physis and extended an additional 0.5 mm. Image reconstruction was performed using NRecon software (Bruker). After any necessary image reorientation, quantitative analysis was conducted with CTAn (Bruker), and 3D image rendering was completed with CTVol.

### 2.3 Wheel access regimen

5, 12, and 24mo old mice (n=6mice/group) were randomly assigned to either a voluntary wheel running or a non-running control group without access to a running wheel. Animals were given a one-week acclimation period in the new cages, followed by a six-week monitored exercise intervention. All mice were individually housed, and running metrics—including elapsed time, total distance, average speed, and maximum speed—were measured daily using a cyclocomputer (CatEye Velo 7). Body mass and food intake were also recorded weekly.

### 2.4 Bone marrow cell extraction, expansion, and differentiation

Bone marrow cells were isolated from the tibiae and femoral (n=9mice/group) via centrifugation as previously described (Amend et al. 2016). Briefly, the tibia and femur were dissected, cleared of all soft tissue, and centrifuged at >12,000 × g to extract the bone marrow. The resulting bone marrow pellet was then resuspended in media and plated in supplemented growth media (alpha-MEM with 20% FBS, 100 U/mL penicillin, and 100 μg/mL streptomycin). After 48 hours, cells were washed with PBS to remove non-adherent cells, followed by the addition of warm media before microscopic observation. Some immune cells remained visible at this stage. Therefore, the media was gently aspirated and slowly dispensed to wash the growth surface. This process was repeated approximately five times to lift some, but not all, of the adherent cells. After aspirating the media, 2 mL of fresh media was added. The cells were then imaged and returned to the incubator to proliferate until reaching approximately 50% confluency. At this confluency, cells were lifted, split onto separate plates to generate sufficient cell numbers for assays, and allowed to proliferate to ∼80% confluency. Cells were then lifted again and plated according to experimental requirements.

### 2.5 RNA-seq analysis

Total RNA was extracted using the RNeasy kit (Qiagen) from three samples per group. The RNA samples were sent to Novogene for mRNA sequencing and analysis. Briefly, the reference genome index was built using Hisat2 v2.0.5, and paired-end clean reads were aligned to the reference genome using Hisat2 v2.0.5. FeatureCounts v1.5.0-p3 was used to count the number of reads mapped to each gene. The fragments per kilobase of transcript per million mapped reads (FPKM) for each gene were then calculated based on gene length and read count. Differential expression analysis was performed using the DESeq2 R package (v1.20.0), which employs statistical routines based on the negative binomial distribution to determine differential expression in digital gene expression data. Resulting p-values were adjusted using the Benjamini-Hochberg method to control the false discovery rate. Genes with an adjusted p-value < 0.05 were considered differentially expressed. These significantly differentially expressed genes were further analyzed using DAVID for pathway enrichment analysis.

### 2.6 Immunofluorescence study of Ki67 and P16

For the immunofluorescence study of Ki67 and P16, cells (n= 3mice/group) at 10k were cultured for 2days in the growth media. After 2days, cells were fixed with 4% formaldehyde and permeabilized by incubation with incubation with 0.3% Triton X-100. Subsequently, cells were incubated in blocking serum prepared in PBS containing 5% Bovine Serum Albumin (Sigma Aldrich, A3912-100G). Primary antibodies Ki67 (Novus, NB110-89717) and P16 (Cell Signaling Technology, 80772S) were incubated on the cells for overnight at 4°C, followed by secondary antibody incubation of Alexa Flour 594 goat anti-rabbit (invitrogen, A11037). For actin staining, cells were incubated with Phalloidin 488 (Cayman Chemical Company, 20549). For nuclear staining, cells were incubated with NucBlue Live Cell Stain (invitrogen, R37605). Primary and secondary concentrations were both 1:300. Imaging was conducted using a Zeiss LSM900 Microscope with 60x/1.4 oil and 20x objective for ki67 and p16 respectively.

### 2.7 Oil Red-O and lipid droplet staining

For adipogenic assays (n=3mice/group), cells were seeded at 150,000 cells per well on 6-well plates and imaging plates for Oil Red O and lipid droplet staining, respectively. Cells were maintained in normal growth media for 24 hours prior to switching to adipogenic media (alpha-MEM with 10% FBS, 100 U/mL penicillin, 100 μg/mL streptomycin, 5 μg/mL insulin, 0.1 μM dexamethasone, and 50 μM indomethacin) for 7 days. The medium was changed three times per week. After differentiation, cells were fixed with 10% formalin for 30 minutes at room temperature for Oil Red O staining as described previously (Kraus et al. 2016). Following fixation, 100% propylene glycol USP (Poly Scientific R&D corporation, s264-8oz) was added for 2 minutes, then replaced with Oil Red O (Poly Scientific R&D corporation, s1848-8oz) to cover the entire monolayer of cells for 15 minutes. After removing the Oil Red O, 85% propylene glycol (Poly Scientific R&D corporation, s264a-8oz) was applied for 1 minute, followed by rinsing the cell cultures with distilled water.

For immunofluorescence analysis of adipogenic differentiation, cells were fixed and stained with LipidSpot 488 (Biotium, 70065) according to the manufacturer protocol. Image analysis and quantification were performed using ImageJ Software 1.52 by calculating the percentage area fraction of fat globules relative to the total area and measuring the mean intensity of fat globules.

### 2.8 Alizarin red and xylenol orange staining

For the osteogenic study (n=3mice/group), cells were seeded at 200,000 cells per well on 6-well plates and imaging plates for Alizarin Red (Sigma Aldrich, A5533-25G) and Xylenol Orange (Sigma Aldrich 398187-1G) assays, respectively. Once the cells reached 100% confluence, the growth medium was replaced with osteogenic medium consisting of alpha-MEM supplemented with 20% FBS, 100 U/mL penicillin, 100 μg/mL streptomycin, 10 mM beta-glycerophosphate, and 50 μg/μL ascorbic acid. The osteogenic medium was refreshed 2–3 times per week for 21 days. After 21 days, cells were fixed with 70% ethanol for 1 hour and stained with 40 mM Alizarin Red for 10 minutes at room temperature to ensure sufficient coverage (Reinholz et al. 2000). After 10 minutes Alizarin Red removed and washed with ddH_2_O for five times. For immunofluorescence analysis of osteogenic nodules, Xylenol Orange staining was performed following a previously established protocol (Wang et al. 2006). Briefly, cells were fixed with 4% paraformaldehyde for 10 minutes at room temperature, washed twice with PBS, and incubated with Xylenol Orange at 4°C overnight. Images were quantitatively analyzed using ImageJ software 1.52.

### 2.9 Real-time PCR

For the analysis of adipogenic and osteogenic gene marker expression, cells were seeded in 6-well plates. RNA was extracted after 7 days (adipogenic differentiation) and 21 days (osteogenic differentiation) using Trizol reagent and the QIAzol RNA extraction kit. Complementary DNA (cDNA) was synthesized using a Script cDNA synthesis kit in a thermal cycler. Primer sequences used in this study are provided below. Real-time PCR was performed using the SSO Advanced system. Data were normalized to 18S rRNA expression, and relative mRNA expression levels were calculated using the 2^−ΔΔCt method. Primers used in the study are given in Supplementary **Table S8.**

### 2.10 Application of low-intensity vibration

Low-intensity vibration (LIV) was applied to aged MSCs (+LIV) at peak magnitudes of 0.7g and 90 Hz, administered four times daily for 20 minutes each session with 1-hour rest intervals at room temperature. Control aged MSCs (−LIV) were removed from the incubator for the same duration and frequency but were not exposed to vibration.

### 2.11 Nuclear morphology and actin fiber measurement

F-actin and nuclear staining were performed on aged MSCs isolated from 5mo, 12mo, and 24mo-old mice. Cells were fixed with 4% paraformaldehyde and permeabilized by incubation with 0.3% Triton X-100. Subsequently, cells were incubated in blocking serum prepared in PBS containing 5% donkey serum (017-000-121, Jackson Immuno Research Laboratories, West Grove, PA, USA). For nuclear staining, cells were incubated with NucBlue Live cell Stain (invitrogen, R37605). For actin staining, cells were incubated with Alexa 488-phalloidin (Life Technologies, Carlsbad, CA, USA). Both primary and secondary antibody concentrations were 1:300.

Our image analysis was divided into three phases: image preprocessing, image segmentation, and post-processing. Z-stack confocal images were analyzed to reconstruct whole-cell actin fibers, perinuclear actin fibers, and nuclei. The method for perinuclear fiber reconstruction as previously described (Nikitina et al. 2024). For whole-cell F-actin reconstruction, the aforementioned method was modified. Initially, images were preprocessed by manually generating binary masks to segment actin and nuclear regions using maximum intensity projections of separate channels. The actin masks were eroded to separate closely spaced cells; after distinct contours were identified from the eroded masks, individual contours were dilated to recover the full cell area. Actin contours were paired with nuclear contours based on maximum overlap. Using these pairs, the 3D confocal image matrix was cropped to extract each nucleus and its associated actin region. To ensure compatibility with the U-Net segmentation model, padding was added as necessary so that dimensions were divisible by 512 pixels. The actin region was then split into 512×512 tiles along the x–y plane. Finally, cross-sectional (x–z) images were generated from both nucleus and actin stacks.

The segmentation and reconstruction protocol for whole-cell fiber reconstruction was performed as previously described by (Nikitina et al. 2024). After segmentation, all 512×512 tiles were stitched together to reconstruct the full segmented 3D mask of the actin region, followed by reconstruction. Nuclear length and width were estimated by fitting an ellipse to the contour based on the nucleus’s maximal projection mask. Height was derived by measuring the maximal height of the reconstructed nucleus, and volume was calculated by summing the segmented area from each slice and multiplying by the resolution. Actin fiber length was estimated by summing the distances between every fifth point along individual fibers in the x–y plane to reduce the zigzag effect of the reconstructed fibers. Fiber volume was computed by summing the cross-sectional areas of each slice and multiplying by the resolution.

### 2.12 Mitochondrial Image analysis and identification of key pathways associated genes

Mitochondrial staining was performed using Tetramethylrhodamine methyl ester (TMRM) on live cells. Cells were incubated in cell culture medium containing TMRM (Invitrogen, T668) and NucBlue Live Cell Stain (invitrogen, R37605) for 15 minutes at 37°C following the protocol from the user guide of the product. After incubation, cells were washed with PBS, replaced with fresh medium, and placed in a temperature- and CO₂-controlled chamber attached to the stage of a confocal microscope. TMRM staining was conducted on aged MSCs isolated from 5mo, 12mo, and 24mo-old mice to compare the effects of aging. Additionally, TMRM staining was performed on aged MSCs after ±LIV treatment, as described earlier, to analyze the mechanical effects on aged MSCs.

Image analysis was performed to quantify TMRM fluorescence. Z-stack images were analyzed to identify mitochondrial fibers, where a binary mask separated the mitochondrial and nuclear regions of cells using separate channels. The mitochondrial masks were eroded to distinguish closely adjacent cells; after distinct contours were identified from the eroded masks, individual contours were dilated to recover the full mitochondrial area. The 3D confocal image matrix was then cropped to extract each nucleus and its associated mitochondrial region. To ensure compatibility with the U-Net segmentation model, padding was added as necessary so that dimensions were divisible by 512 pixels. The mitochondrial region was subsequently split into 512×512 tiles along the x–y plane. Cross-sectional (x–z) images were generated from both the nucleus and the mitochondria stacks. After segmentation, all 512×512 tiles were stitched together to reconstruct the full segmented 3D mask of the mitochondrial region.

Mitochondrial fiber length was estimated by summing the distances between every fifth point along individual fibers in the x–y plane to reduce the zigzag effect of the reconstructed fibers. Fiber volume was calculated by summing the cross-sectional areas of each slice and multiplying by the resolution.

We used data set (http://software.broadinstitute.org/gsea/msigdb) from the molecular signature database (MSigDB) for mitochondrial fission, fusion, mTOR, genome maintenance and oxidative phosphorylation. GSEA analysis was performed for all genes with set NOM. *p*<0.05 and thresholds of size >20. Heatmap of gene expression for each pathway were presented in supplementary figures S6 through S10. Top 5 up and down-regulated genes from each pathway were presented in the Figure 7.

### 2.13 Atomic Force Microscopy of Cells Viscoelastic Properties

Aged MSCs were seeded at a density of 4,000 cm^2^ to ensure isolated single cell engagement. Atomic force microscopy (AFM) imaging was conducted after 24 hours of culture and completed prior to 48 hours in alpha-MEM supplemented with 20% FBS, 100 U/mL penicillin, 100 μg/mL streptomycin. To minimize mechanical alterations from cell death under ambient conditions, each plate was imaged for no longer than 1 hour.

AFM was performed using a NanoWizard V Bioscience (Bruker, Billerica, MA, USA) integrated with a Zeiss LSM Airyscan (Zeiss, Oberkochen, Germany) for cell visualization. Mechanical measurements were acquired with MLCT-SPH-1UM probes (Bruker, Billerica, MA, USA), and viscoelastic data were obtained using MicroRheology Force Mapping in the JPK SPM software. Force ramps were segmented into four phases: (1) an extension period to a setpoint of 0.6 nN, (2) a 500 ms constant-height hold, (3) a modulation phase consisting of 20 cycles at 10 Hz with a 2 nm amplitude, and (4) a retraction period. All segments were sampled at 2048 Hz. 8 x 8 pixel force maps were acquired either over entire cells or within a 100 × 100 µm region centered on the nucleus. Mechanical data were processed using JPK Data Processing software. Raw force curves were first smoothed using a Gaussian filter with a width of 3.00, then baseline-corrected over a standardized fit range of 50–100% of the x-axis (height) and further corrected for offset and tilt. Contact point correction was applied to define the contact point as the first crossing of the y-axis (vertical deflection). Vertical tip position (piezo height minus cantilever deflection) was used to account for cantilever bending. Apparent Young’s modulus was obtained by fitting standard force curves with a Hertz/Sneddon model assuming a spherical indenter geometry, with a nominal probe radius of 1 µm. The modulation segment was analyzed using vertical tip position as the indentation channel and vertical deflection as the force channel, with a standardized fit range of 10–90% of the modulation segment. Dynamic Young’s modulus, storage modulus, loss modulus, loss tangent, shear storage modulus, and shear loss modulus were all derived from the modulation segment, also under the assumption of a spherical indenter geometry. For both static and dynamic fits, the Poisson’s ratio was fixed at 0.40 (Sen et al. 2022).

Apparent Young’s modulus values were restricted to a physiological range of 0–30 kPa to exclude spurious contacts with artifacts or the substrate, and outliers were removed using a 1.5× interquartile range (IQR) criterion. Dynamic mechanical parameters were filtered using the same criteria applied to the Dynamic Young’s modulus values. In addition, dynamic datasets were further restricted by excluding points with a loss tangent greater than 1 to eliminate false engagements in the fluid environment.

Nuclear and cytoplasmic mechanical data were extracted by overlaying brightfield cell images with height maps. Pixels corresponding to increased height were used to identify and confirm engagements over the nuclear region, which were then isolated using a custom script. All remaining data were assigned to the cytoplasmic region.

### 2.14 Statistical Analysis

All statistical analyses were conducted using GraphPad Prism software (version 10.2.0; GraphPad Software LLC, San Diego, CA, USA). The study primarily involved comparisons among three distinct age groups. For multiple group comparisons, one-way analysis of variance (ANOVA) was employed. Pairwise comparisons between two groups were performed using unpaired Student’s *t*-tests. Data are presented as mean ± standard deviation (SD), and a *p*-value of less than 0.05 was considered statistically significant.

## 3 Results

### 3.1 Bone quality and exercise performance decreased with chronological aging

Tibiae were harvested from one leg of mice aged 5, 12, and 24 months and prepared for micro-ct analysis (**Figure 1a**). The micro-CT data included measurements of bone volume to total volume ratio (BV/TV), trabecular number, trabecular spacing, cortical area to total area ratio (Ct.Ar/Tt.Ar), cortical area, and cortical thickness. Results demonstrated a significant decrease in BV/TV with age; compared to 5-month-old mice, BV/TV decreased by 41.52% (p < 0.01) in 12-month-old and 41.11% (p < 0.01) in 24-month-old mice (**Figure 1b**). Trabecular number also significantly declined with age, decreased 45.85% (p < 0.001) and 43.17% (p < 0.01) in 12mo and 24mo-old mice, respectively (**Figure 1c**). Conversely, trabecular spacing increased significantly by 50.96% (p < 0.0001) in 12mo and 93.44% (p < 0.0001) in 24mo mice compared to 5 mo, with an additional 28.14% increase from 12mo to 24mo (p < 0.0001) (**Figure 1d**). The cortical area to total area ratio decreased by 6.46% (p < 0.05) and 24.89% (p < 0.0001) in 12mo and 24mo-old mice, respectively, relative to 5mo-old mice (**Figure 1e**). Furthermore, a 19.70% (p < 0.0001) cortical area reduction was observed in 24mo mice compared to 12mo mice (**Figure 1e**). Cortical area also declined by 7.8% (p < 0.05) and 28.36% (p < 0.0001) in 12mo and 24mo-old groups relative to 5mo, with a further 22.17% decrease in 24mo-old mice compared to 12mo (p < 0.0001) (**Figure 1f**). Cortical thickness significantly decreased by 3.9% in 12mo mice and 27.28% (p < 0.0001) in 24mo mice compared to 5mo, with a 24.23% reduction in 24mo relative to 12mo (p < 0.0001) (**Figure 1g**).

**FIGURE 1.**
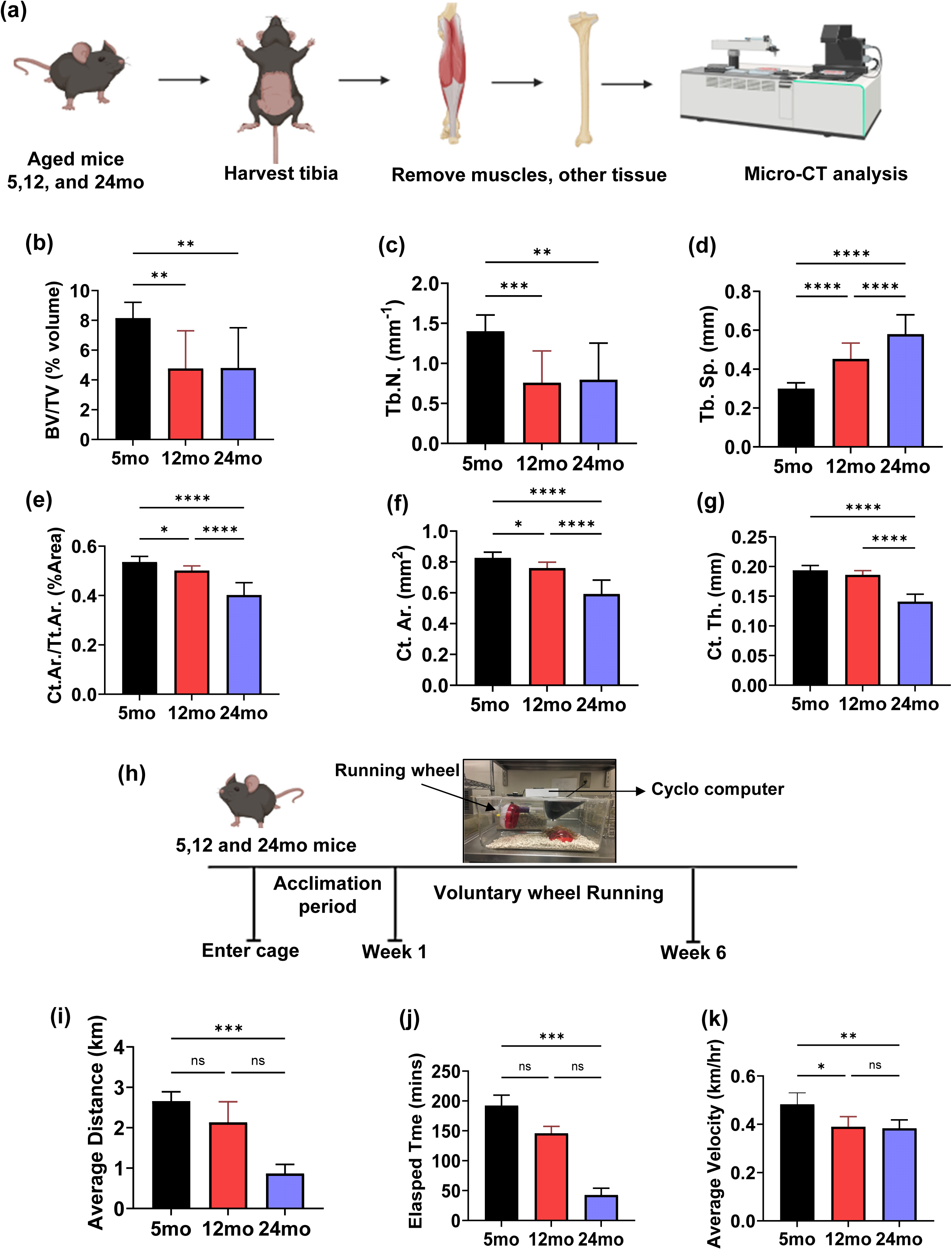
Bone microarchitecture and exercise performance significantly decrease with aging. (a) Bone extracted from the 5,12 and 24 mo aged mice and analysis in micro-Ct. Trabecular measurements include (b) bone volume fraction (BV/TV), (c) trabecular number (Tb.N.), (d) trabecular spacing (Tb.sp.). Cortical measurements include (e) total cross-sectional area inside the periosteal envelope(Tt.Ar.) cortical area fraction (Ct.Ar./Tt.Ar.),, (f) cortical bone area (Ct.Ar.), and (g) average cortical thickness (Ct.Th.) (h) Exercise regimen timeline schematic. Running metrics tracked by cyclometer measuring (i) average distance, (j) elapsed time, and (k) average velocity. *p<0.05, **p<0.01, ***p<0.001, ****<0.0001

Another group of mice was given voluntary access to a running wheel or no wheel and allowed a one-week acclimation period and voluntary running metrics were tracked for six-weeks (**Figure 1h**). Outcomes included food intake, body mass as well as daily averages of distance, elapsed time, and velocity. Body mass and food intake followed similar trends and stayed the same across all age groups (**Figure S1a,1b**). Notably, the average running distance decreased with age: compared to 5mo-old mice, 12mo and 24mo-old mice exhibited reductions of 19.62% and 67.30% (p < 0.001), respectively (**Figure 1i**). Elapsed time also decreased significantly with age, showing declines of 24.08% and 77.71% (p < 0.001) for 12mo and 24mo groups, respectively (**Figure 1j**). Similarly, average running velocity was reduced by 19.23% (p < 0.05) and 20.61% (p < 0.01) in 12mo and 24mo-old mice compared to 5mo-old mice (**Figure 1k**).

### 3.2 RNA seq analysis identifies key factors involved in MSCs senescence with chronological aging

Both femur and tibias from the contralateral legs (n=11/grp) were used for cell extraction and experiments were plated for RNA-seq, adipogenic and osteogenic differentiation (imaging and qPCR), immunostaining (p16, Ki67), F-actin analysis, mitochondria analysis and atomic force microscopy (**Figure S1c**).

Our study investigated RNA-seq data and gene expression changes between the 5mo, 12mo, and 24mo samples (**Figure 2a-2f, Figure S2** n= 3/group). Supplementary Figure 2 showed a volcano plot for the comparison of gene expression level between two age groups based on their fold change and significance levels where the gray and green data points for *p* > 0.05, blue points for *p* < 0.05 and red points for significantly upregulated (right side) and downregulated (left side) genes. A total of 2,369 genes were found to be significantly differentially expressed between the 5mo and 12mo samples; among these, 108 genes were upregulated (Log_2_FC > 1), while 419 genes were downregulated (Log_2_FC < -1, **Figure S2a**). Between the 12mo and 24mo samples, 1,569 genes showed significant differential expression, including 644 upregulated (Log_2_FC > 1) and 129 downregulated (Log_2_FC < -1) genes (**Figure S2c**). Similarly, the comparison between the 5mo and 24mo samples revealed 945 significantly differentially expressed genes, with 283 upregulated (Log_2_FC > 1) and 176 downregulated (Log_2_FC < -1) (**Figure S2e**).

**FIGURE 2.**
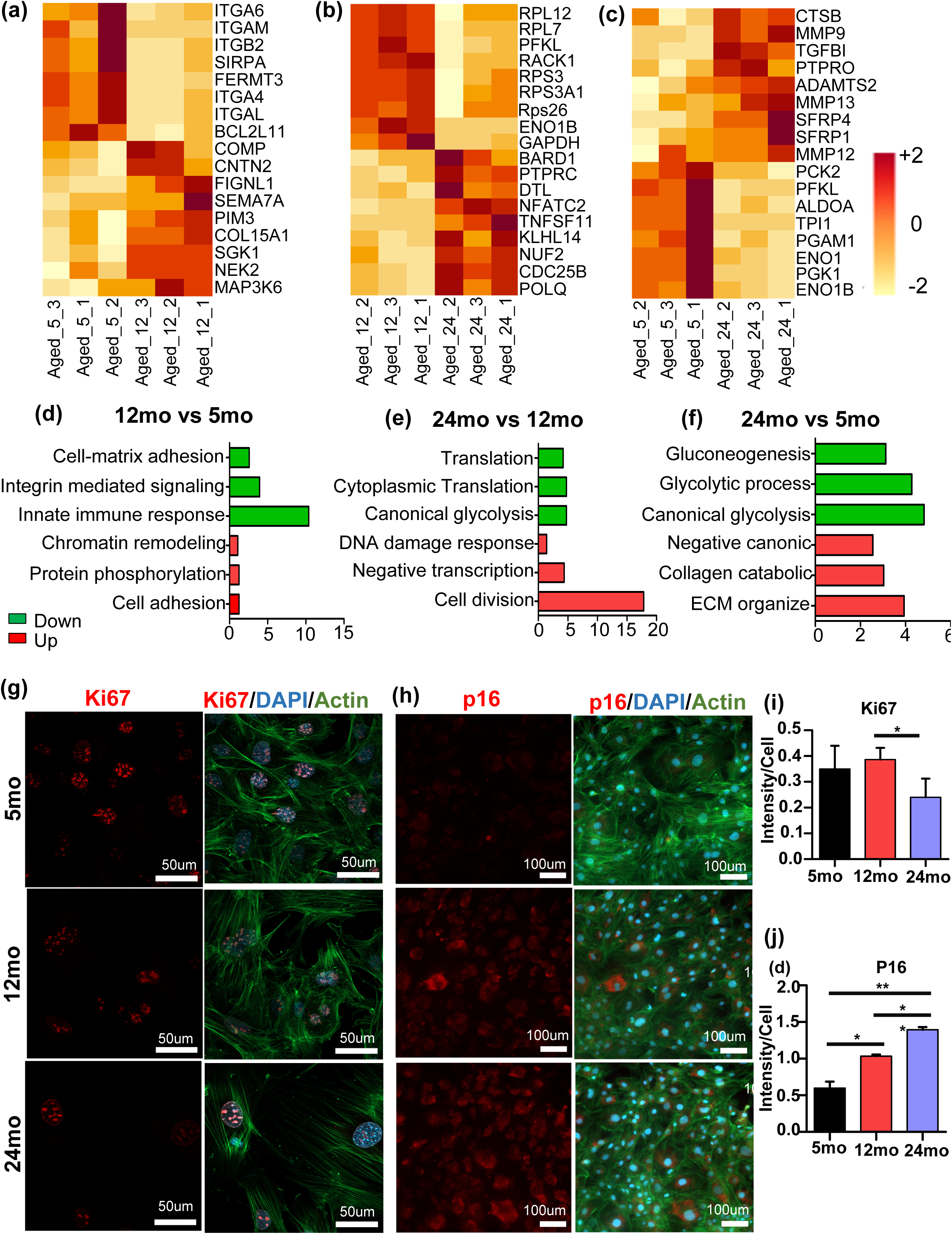
Dynamic changes in gene expression profiles and decrease the proliferation marker Ki67 and increase the senescence marker p16 of MSCs in chronological aging. (a,b,c) The GO terms associated top 3 highly upregulated and downregulated gene responsible for pathways were represented as a heatmap. (d,e,f) Depicts the 3 significantly enriched GO terms associated with upregulated and downregulated gens,as identified by DAVID. The x-axis indicates the log Benjamin score for each GO term. They y-axis represent the detailed classification of the GO terms. GO terms associated with upregulated and downregulated are highlighted with red and green respectively. Immunofluorescence staining of (g) Ki67 and (h) p16 and their expression for 5mo,12mo,and 24mo MSCs. Quantitively analysis and intensity measurement of confocal images of (i) Ki67 and (j) P16 for 5mo, 12mo, and 24mo of MSCs. Quantitative measurements were presented as an average ± SD and significance level determined via one-way ANOVA among three groups.*p<0.05, **p<0.01, ***p<0.001, ****p<0.0001.

Gene ontology analysis using DAVID was conducted to elucidate the biological significance of these differentially expressed genes. Heatmaps of key gene expression levels were illustrated in **Figure 2a, 2b, and 2c** for 12mo vs 5mo, 24mo vs 12mo, and 24mo vs 5mo, respectively. Based on the Benjamini adjusted significance level, three key biological processes relevant to aging were identified. **Figure 2d, 2e, and 2f** depict the biological processes associated with upregulated and downregulated genes in the 12mo vs 5mo, 24mo vs 12mo, and 24mo vs 5mo comparisons, respectively. The results indicated that 12mo MSCs, when compared to 5mo samples, showed a downregulation of in genes related to cell-matrix adhesion, integrin-mediated signaling, and innate immune response (**Figure 2d**). For the 24mo vs 12mo comparison, upregulated genes were associated with cell division, negative regulation of transcription, and DNA damage response, whereas downregulated genes were mainly involved in translation and canonical glycolysis (**Figure 2e**). Finally, 24mo samples, when compared to 5mo, upregulated genes were implicated in the negative regulation of Wnt signaling and collagen catabolic processes. Downregulated genes were involved in glycolysis and gluconeogenesis (**Figure 2f**). Data for each differentially regulated pathway is presented in supplementary tables S2-S7.

### 3.3 Proliferation Markers Ki67 decreased and senescence markers p16 increased with aging

To assess the proliferative capacity of aging primary MSCs, we utilized the Ki67 antibody, a well-established marker of cellular proliferation (Scholzen & Geredes 2000). Immunofluorescence imaging revealed Ki67 expression across three different age groups (**Figure 2g**). Quantitative analysis demonstrated that Ki67 intensity/cell was significantly reduced by 37.93% (p<0.05) in 24mo cells compared to 12mo cells, however no significant changes between 5mo to 24mo (**Figure 2i**).

We next quantified P16, a widely recognized as a biomarker of cell cycle arrest associated with senescence (Safwan et al. 2022). Immunofluorescence analysis indicated an age-dependent increase in P16 intensity/cell (**Figure 2h**). Quantitative intensity measurements showed that P16 intensity/cell increased by 57.13% (p<0.05) and 42.22% (p<0.01) in 12mo and 24mo, respectively, when compared to 5mo (**Figure 2j**).

### 3.4 Aging increases the adipogenic and decreases the osteogenic differentiation of MSCs

In **Figure 3a**, Oil Red O staining was evident in all groups, with quantitative analysis revealing an age-dependent increase in fat droplet accumulation. Specifically, the area fraction analysis demonstrated that fat globule content in 24mo cells increased by 36.91% (p<0.05) and 24.33% compared to the 5mo and 12mo groups, respectively (**Figure 3c**). Immunofluorescence imaging using a Lipid Spot 488 staining further confirmed enhanced adipogenesis with aging (**Figure 3d**).

**FIGURE 3.**
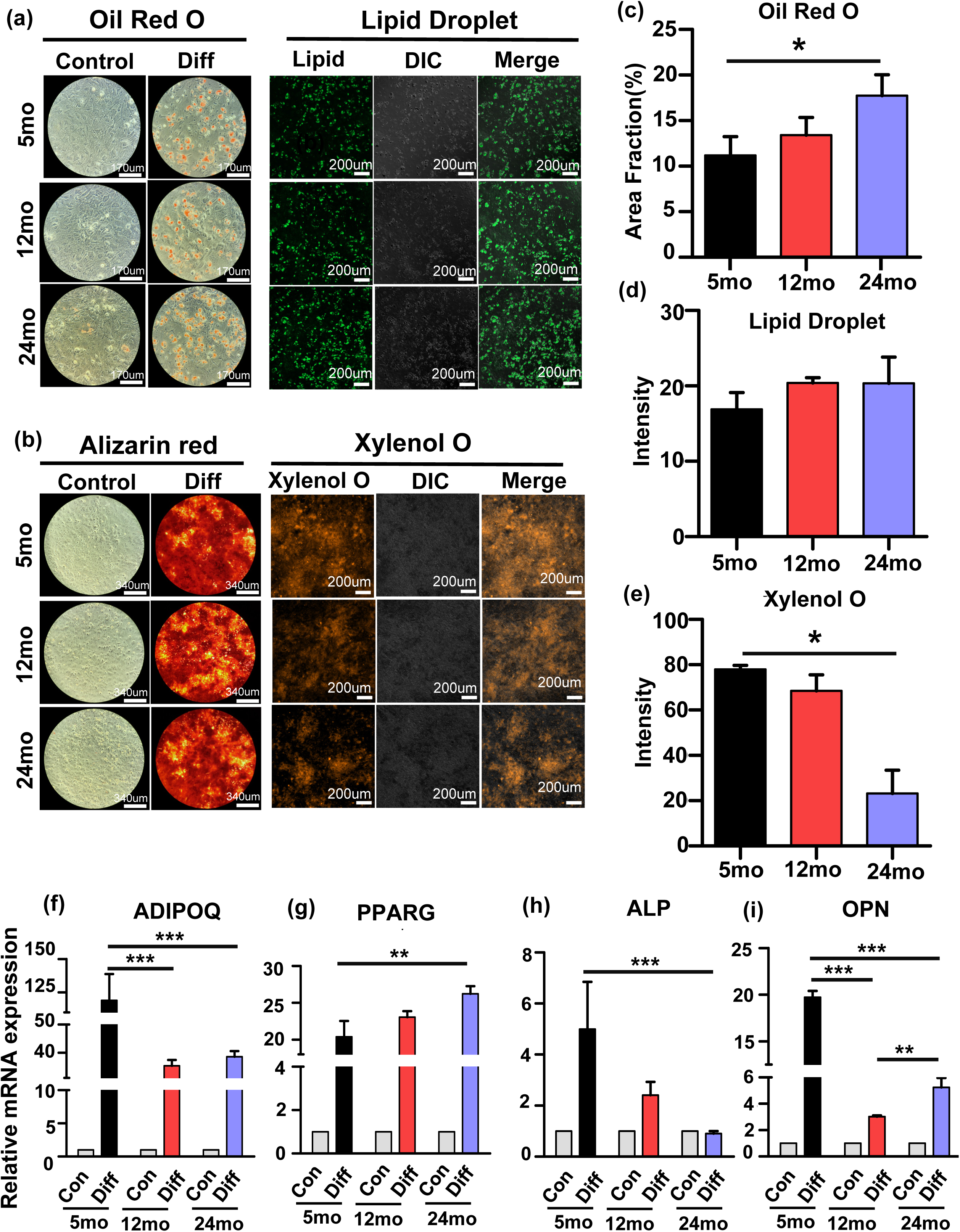
MSCs displays increase adipogenesis and decrease osteogenesis in chronological aging. (a) Oil O red and lipid droplet staining of fat globule in normal growth media and adipogenic media and (b) Alizarin red and Xylenol O staining represent the nodule formation after osteogenic differentiation for 5mo, 12mo, and 24mo MSCs. (c) Quantitative analysis of the area fraction of fat globule from Oil O red staining. (d) Mean intensity of Lipid droplet from the immunofluorescence images. (e) Mean intensity of Xylenol O for 5mo, 12mo, and 24mo MSCs. qPCR data showed (f,g) adipogenic markers Adipoq, PPRG and (h,i) osteogenic markers APL,OPN expression level among the three groups. Result presented as an average ± SD and significance level determined via one-way ANOVA.*p<0.05, **p<0.01, ***p<0.001, ****p<0.0001

Osteogenic nodule formation was evaluated through Alizarin Red and Xylenol Orange staining, followed by image intensity quantification (**Figure 3b**). Immunofluorescence analysis of Xylenol Orange revealed a significant reduction in staining intensity in the 24mo group (**Figure 3e**). Quantitative assessment demonstrated that nodule formation in 24mo cells decreased by 70.25% (p<0.05) and 66.11% compared to 5mo and 12mo samples, respectively.

Gene expression analysis via qRT-PCR was conducted to compare adipogenic and osteogenic markers across the three age groups. After 7 days of adipogenic induction, adipogenic gene ADIPOQ showed a significant decrease in 12mo (70%, p<0.001) and 24mo (68%, p<0.001) samples when compared to 5mo samples (**Figure 3f**), indicating more advanced adipogenic stage. When we compared the PPARG expression, 24mo cells were higher compared to 5mo (**Figure 3g**, 30%, p<0.01).

Following 21 days of osteogenic culture, mRNA levels of osteogenic markers ALP and OPN were measured. ALP expression was highest in the 5mo samples, with significant reductions observed in aged groups with 24mo decreasing 81% (p<0.001, **Figure 3h**). Similarly, OPN expression significantly decreased in 12mo (84%, p<0.001) and 24mo (73%, p<0.01) cells compared to 5mo cells (**Figure 3i**).

### 3.5 Chronological aging induces Perinuclear actin Loss, nuclear flattening, and cellular stiffening

As RNA-seq analysis indicated possible changes in the integrin signaling, we next evaluated the F-actin and nuclear structure. **Figure 4a** presents the maximum projection confocal images and F-actin reconstructions using our U-net based machine learning algorithm (Nikitina el al. 2024) F-actin measurements include fiber length, volume, and number for both the entire cell (**Figure S3a-c**) and the perinuclear area (**Figure 4b-d**). Nucleus measurements (**Figure 4e-g**) include volume, length and projected area. Total cell F-actin quantification results indicated no significant differences except 12mo cells showing a slight decrease in average fiber volume compared to 5mo (21.30%, p<0.01). Comparing nuclei-associated fibers we found a larger decrease in fiber volume in both 12mo and 24mo cells with 12mo being 21.32% lower (p < 0.01) compared to 5mo.

**FIGURE 4.**
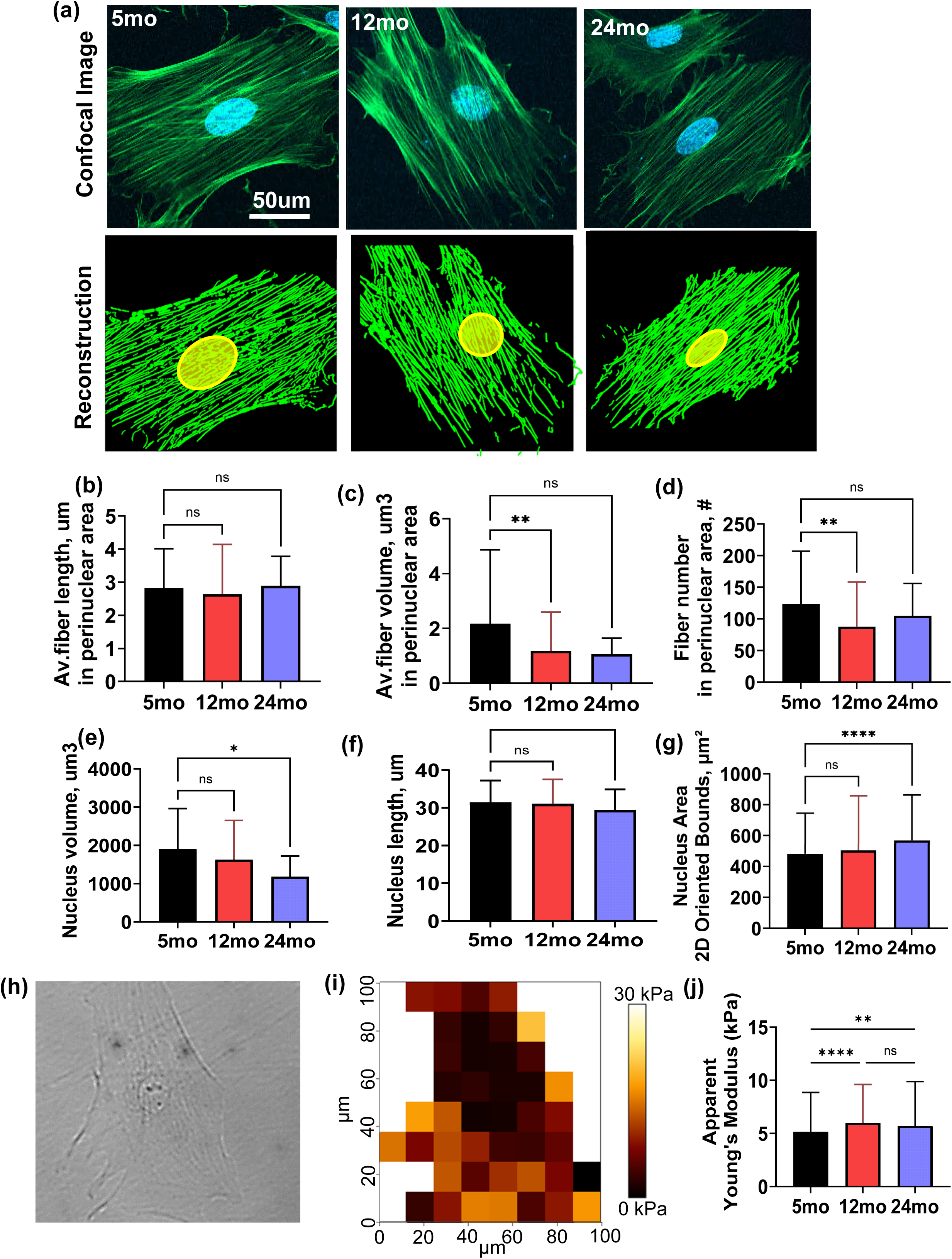
Age associated changes of perinuclear actin fiber, nucleus morphology and viscoelastic properties. (a) Representative fluorescence microscopy and their reconstruction images display the actin fiber in green and nuclei in blue. Quantitative measurements include (a) avg. fiber length, (c) number and (d) volume in prenuclear area were presented. (e) Nucleus volume (f) length (g) area were measured and presented as an as an average ± SD. (h) Example of brightfield image of isolated single cell with (i) force map overlay representing dynamic Young’s modulus (kPa). (j) Significant increases in apparent Young’s modulus were observed between 5mo and 12mo, and 5mo and 24mo MSCs (p < 0.0001 and p = 0.006, respectively), with no significant difference between 12mo and 24mo. Significance level determined via one-way ANOVA among three groups.*p<0.05, **p<0.01, ***p<0.001, ****p<0.0001.

Similarly, fiber number in perinuclear area also showed a downward trend in both 12mo and 24mo cells with only 12mo being significantly reduced (28.96%, p<0.01) compared to 5mo. With observed changes in the nuclei-associated fibers, nuclear measurements showed 38.06% decreased volume (p<0.05) despite increasing area projection (17.99%, p<0.0001).

As image analysis showed structural changes with age, we next examined potential age-related changes in MSC viscoelastic properties using AFM MicroRheology Force Mapping of isolated single cells. Shown in **Figure 4h–j**, the probe was able to engage multiple points across the cell, which indicated that measurements on top of nuclei was in general softer than the cytoskeleton region. Averaged across the cell, MSCs showed a significant increase in apparent Young’s modulus between 5mo and 12mo (16.11%, p<0.0001) as well as 5mo and 24mo (10.61%, p<0.01), respectively), with no significant differences between the 12mo and 24mo groups (**Figure 4j**). Similar trends were observed for other elastic measurements, including dynamic Young’s modulus (**Figure S3d** 12mo vs 5mo, 9.2%, p<0.05 and 24mo vs 5mo, 8.4%, p<0.05), storage modulus (**Figure S3e** 12mo vs 5mo, 9.5%, p<0.05 and 24mo vs 5momo, 8.8%, p<0.05), and shear storage modulus (**Figure S3h**, 12mo vs 5mo, 9.57%, p<0.05 and 24mo vs 5mo, 8.88%, p<0.05). No significant differences were found in viscoelastic measurements between age groups, including loss modulus and shear loss modulus (**Figure S3f, S3i**). Consequently, the loss tangent was significantly decreased between 5mo and 12mo (9.5%, p<0.0001), as well as 5mo and 24mo (9.7%, p<0.0001), with no change between 12mo and 24mo, reflecting an increased dominance of elastic behavior relative to viscous behavior in older cells (**Figure S3g**). Overall, these findings indicate that aged MSCs are mechanically stiffer and exhibit more elastic, less viscous behavior compared to younger cells.

### 3.6 Age-related mitochondrial dysregulation attenuates mitochondrial response to mechanical stress

Mitochondria is essential for maintaining cellular health and homeostasis. To investigate the effects of aging on MSCs, we performed TMRM dye staining on MSCs derived from three different age groups (**Figure 5a**). The results were quantified in terms of average mitochondrial length, average fiber volume, and mitochondrial number per cell. Compared to 5mo MSCs, the 12mo and 24mo groups exhibited significant increases in mitochondrial length by 17.14% (p < 0.05) and 10.60% (p < 0.05), respectively (**Figure 5b**). Similarly, mitochondrial volume increased by 25.97% (p < 0.05) in 12mo MSCs and by 9.86% in 24mo MSCs relative to the 5mo group (**Figure 5c**). No significant differences were observed in mitochondrial number across the three age groups (**Figure 5d**) while there was a trend toward decrease with increasing age. In addition, average mitochondrial intensity showed decreased intensity from 5mo to 12mo was 8.3% (p<0.05) and increased from 5mo to 24mo 0.8% (p<0.01), and from 12mo to 24mo 9.85% (p<0.0001) **(Figure S5b**). These structural changes were accompanied by coordinated transcriptional remodeling across mitochondrial fusion, fission, oxidative phosphorylation, mTOR signaling, and genome-maintenance MSigDB gene sets when comparing 12- and 24mo old MSCs to 5mo controls.

**FIGURE 5.**
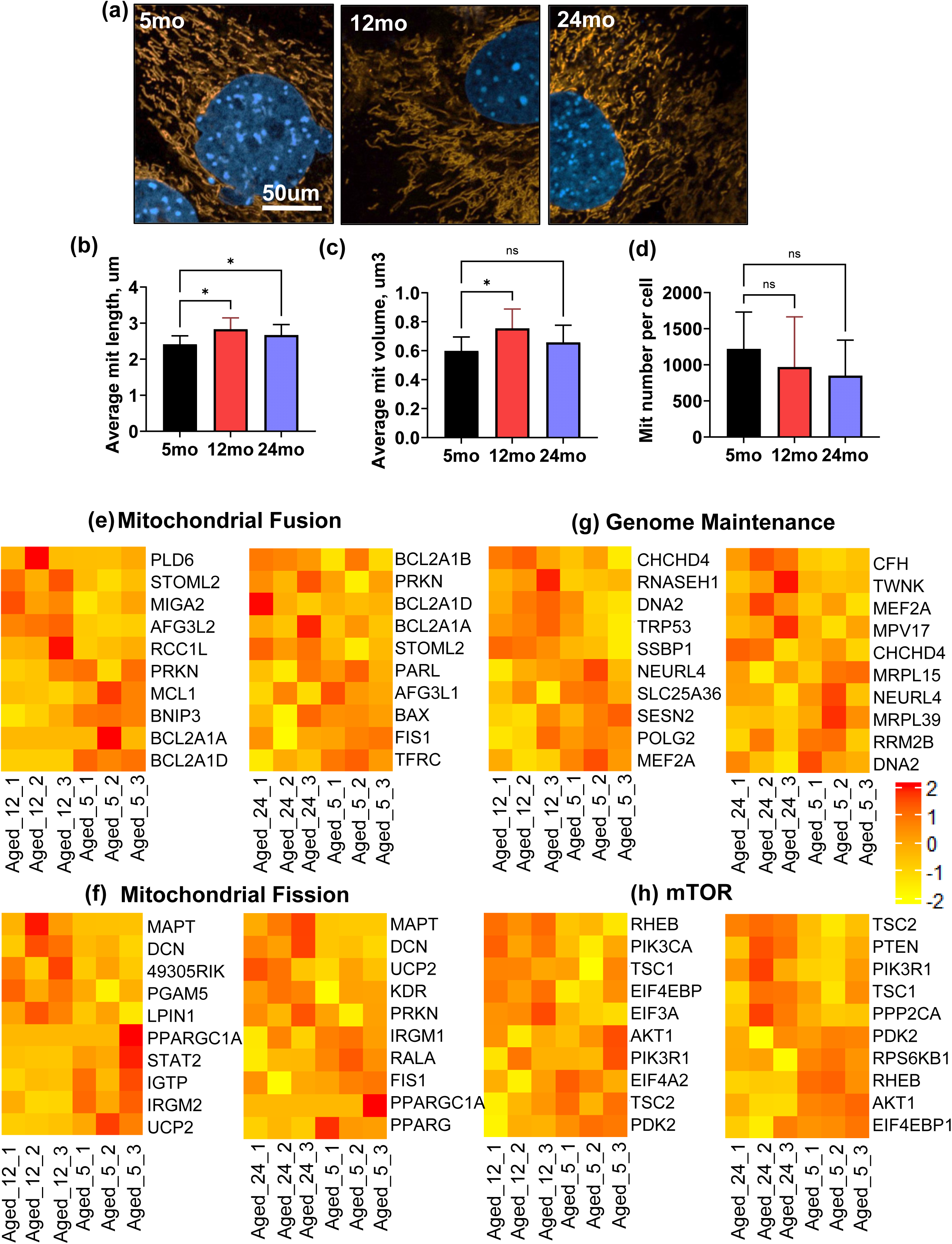
Aging increase the mitochondrial fiber length and induce inflammation signaling pathways. (a) Fluorescence images represent the mitochondrial fiber in orange and nuclei in blue color. Quantitative measurement represent (b) mitochondrial length (c) mitochondrial volume and (d) mitochondrial number per cells. Heatmap images showed the GSEA analysis of the different types of pathways and top 5 upregulated and downregulated genes during comparison of 12mo vs 5mo and 24mo vs 5mo. The pathways were (e) Mitochondrial fusion (f) Mitochondrial Fission (g) genome maintenance and (h) mTOR Results represent as an average ± SD. Significance level determined via one-way ANOVA among three groups.*p<0.05, **p<0.01, ***p<0.001, ****p<0.0001.

Using the mitochondrial fusion gene set (**Figures 5e and S7**), 12mo MSCs exhibited a predominantly increased fusion-associated signature, with upregulation of membrane biogenesis and remodeling genes such as PLD6 (Watanabe et al. 2011), STOML2 (Christie et al. 2011), and MIGA2 (Hong et al. 2022), whereas 24-month MSCs showed a more balanced pattern with additional modulation of apoptosis and mitophagy regulators, including PRKN (Narendra et al. 2008) and BCL2A1 genes (Vogler, 2012). For mitochondrial fission (**Figures 5f and S8**), both 12- and 24-month MSCs displayed a balanced number of up- and downregulated genes. We observed and age-dependent increase in MAPT (Pérez et al. 2018), a well-known mitochondrial disruptor in neurodegeneration, and increased expression of DCN, an ECM protein and mitochondrial regulator (Neil et al. 2022), as well as the stress-responsive factor UCP2 (Brand & Esteves, 2005). In contrast, expression of core fission machinery components decreased, including FIS1 (Serasinghe & Yoon et al. 2008). mTOR-pathway genes (**Figures 5h, S10**) including RHEB (Parmar & Tamanoi, 2010), PIK3CA (Gao et al. 2011), TSC1 (Huang & Manning, 2008), and EIF4EBP1 (Pause et al. 1994) shown an increase at 12mo suggesting of activation of mTORC1 and translational control while 24mo showed an increase in negative or modulatory regulators including, TSC2, PTEN (Feng et al. 2021), PPP2CA (Chen et al. 2022) and reduced AKT1 (Xie et al. 2022) and EIF4EBP1 (Pause et al. 1994). Genome-maintenance gene sets (**Figures 5g, S9**) at 12mo showed an upregulation of mtDNA replication, repair, and processing factors including RNASEH1 (Cerritelli et al. 2003), DNA2 (Zheng et al. 2008), SSBP1 (Jiang et al. 2021), POLG (Chan et al. 2009). While 24mo MSCs showed decreases in mitochondrial ribosomal and repair components including MRPL15 & 39 (Cheong et al. 2020), RRM2B (Awoyomi et al. 2025) and DNA2, consistent with dynamic adaptation of mitochondrial genome integrity pathways during MSC aging. Oxidative phosphrylation-related transcripts (**Figures S6, S 11&12**) were broadly increased at 12 months and more heterogeneously altered at 24 months. Upregulated gene at 12mo **(Figures S6)** involved mitochondrial protein catabolic process AFG1 (Germany et al. 2018) and response to oxidative stress COQ7 (Laredj et al. 2014). On the other hand, at 24mo **(Figures S6)** upregulated genes ACNT3 and TNF were related to negative regulation of oxidative phosphorylation (Berman & North, 2010), and the downregulated genes including COX7B2, DGUOK, MT-ND4L, NDUFAB1 responsible for reducing the ATP synthesis (Guzman et al. 2026)

Since our results indicated changes in mitochondrial stress, we next examined whether aged MSCs mitochondria can respond to mechanical stress by assessing changes in mitochondrial morphology following a 72- hour low-intensity vibration (LIV) treatment (**Figure 6a**). MSCs from 5mo old mice showed a robust increase in average mitochondrial length (26.32%, **Figure 6d**, p < 0.0001) and volume by (35.45%, **Figure 6e**, p < 0.001), as well as a 44.70% decrease in mitochondria number (**Figure 6f**, p < 0.01) when compared to untreated controls (-LIV). MSCs from 12mo and 24mo old mice exhibited smaller increases in response to LIV with mitochondrial length only increasing half of 5mo in 12mo samples (**Figure 6i**, 13.62%,) (p<0.01) and even less in 24mo samples (10.06%, **Figure 6n**, p < 0.05). Neither mitochondrial volume nor mitochondria number showed any significant change in these older groups. Moreover, the intensity measurement data showed LIV decreased the intensity for 5mo (**Figure S5f**, 41%, p<0.0001) and 12mo (**Figure S5i,** 16.53%, p<0.0001) and increased for 24mo (**Figure S5l,** 20.5%, p<0.0001).

**FIGURE 6.**
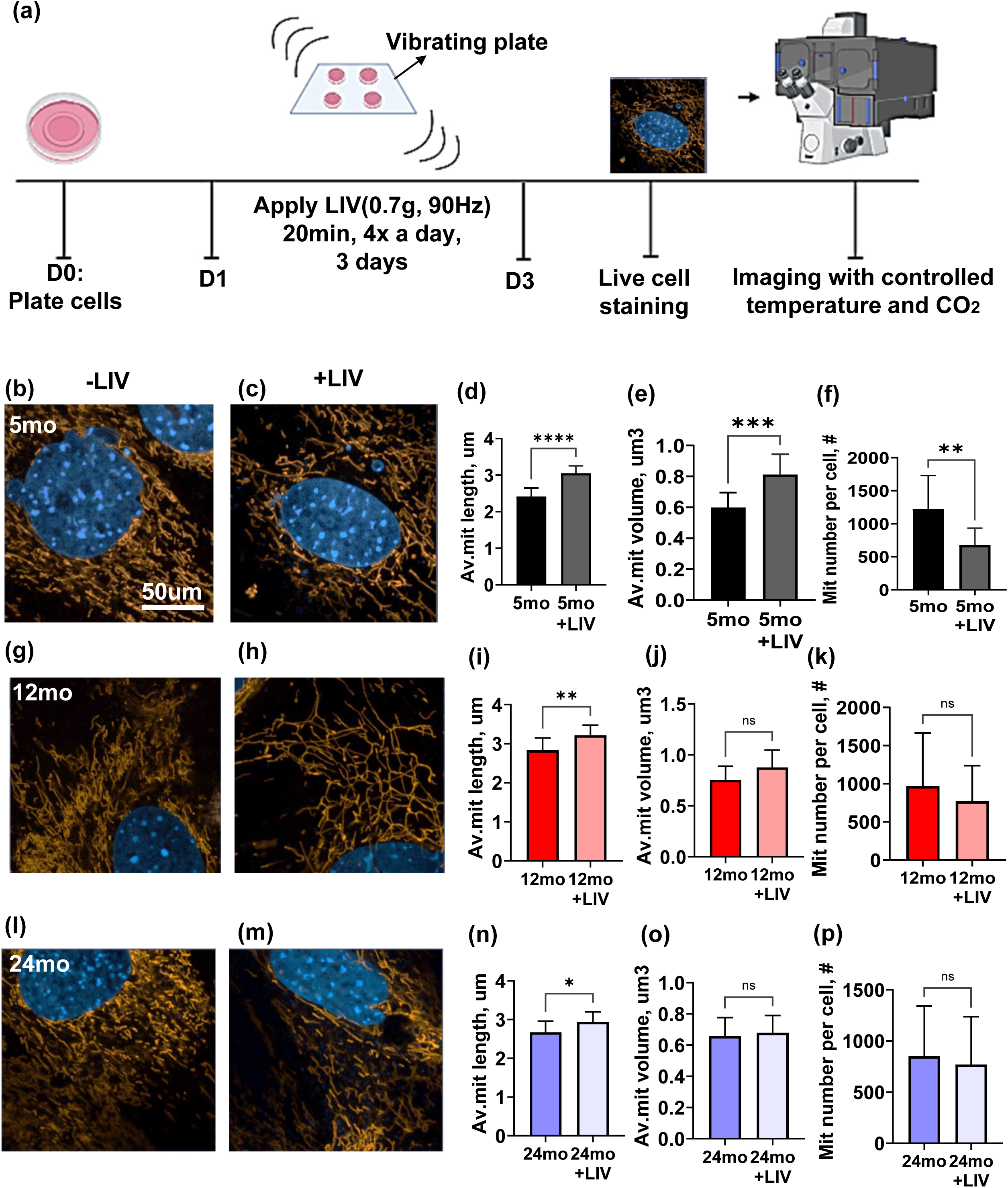
Low intensity vibration stress response in mitochondria decline with aging. (a) Diagram of experimental workflow for LIV application on MSCs. Images showed the mitochondria fiber in orange and nucleus in blue color for -LIV or +LIV treatment of 5mo (b, c), 12mo (g, h) and 24mo (l, m). Comparison of –LIV, +LIV treatment showed quantitively measurement of avg. mito length, avg. mito volume and mito number per cells. Bar graph represnt (d) Avg mito length, (e) avg mito volume (f) mito number per cell for 5mo, (i) Avg mito length, (j) avg mito volume (k) mito number per cell for 12mo, and (n) Avg mito length, (o) avg mito volume (p) mito number per cell for 24mo, between -LIV and +LIV treatment. Results represent as an average ± SD. Significance level determined via student t-test.*p<0.05, **p<0.01, ***p<0.001, ****p<0.0001.

## 4 Discussion

MSCs are a multipotent cell population capable of differentiating into tissue-specific lineages such as adipocytes, chondrocytes, and osteoblasts (Han et al. 2019, Zhang Y et al. 2012). These cells play a crucial role in maintaining biological homeostasis and contributing to the development and regeneration of skeletal tissues. Although it is well-established that MSC activity declines with age, the specific effects of aging gene expression profiles and subcellular structure remain variable from assay to assay and thus requires repeatable methods that can capture aspect of aging to advance further investigation. In this paper, using a previously described method to extract stromal cell populations, we have developed and *in vitro* assay system that extracts MSCs from animals with varying ages (Birks et al. 2024). Findings suggest that we can delineate age-related changes in MSCs *in vitro* which will be useful for future studies to further investigate our findings. Most notably, our method successfully captures previously established outcomes of increased adipogenic bias and decreased proliferative capacity and increased senescence. First volumetric F-actin and mitochondria segmentation along with full-field dynamic mechanical measurements, we have found that despite small changes in F-actin cytoskeleton, cells stiffen, and mitochondria shows increasing dysfunctional markers with increasing age, ultimately resulting in diminished ability to respond mechanical stress.

At first, we investigated the expression of MSC-specific markers gene from RNA-seq data of 5mo, 12mo, and 24mo groups. Based on the list of commercially available RT^2^ profiler^TM^ PCR array mouse mesenchymal stem cells (GeneGlobe ID: PAMM-082zf-2|Cat. No.: 330231| RT^2^ Profiler PCR Arrays), we found out of 85 genes, 68 genes (12mo vs. 5mo) and 67 genes (24mo vs. 5mo) showed no significant expression changes. These findings suggest that cells from all age groups largely retained their progenitor-like characteristics. Fold-change values for all MSC markers are summarized in **Table S1**. We further analyzed 27 MSC differentiation-related genes, including markers of osteogenesis (11 genes), adipogenesis (3 genes), chondrogenesis (12 genes), myogenesis (2 genes), and tenogenesis (4 genes). Based on a ≥2-fold change threshold, no adipogenic markers were differentially expressed, from 5mo to 12mo, one osteogenic gene (BMP6) and one chondrogenic gene (ABCB1a) were upregulated ≥2-fold. In the 24mo vs 5mo comparison, only two osteogenic genes (HNF1A and BMP6) and two chondrogenic genes (LTGX and ABCB1A) showed ≥2-fold increases in expression. These results indicate minimal age-related shifts toward specific differentiated lineages. Further as shown in **Table S1**, PTPRC and IGF1 expression levels decreased by approximately two-fold in 12mo relative to 5mo samples. PTPRC is associated with age-related immune decline, while IGF1 plays a key role in cell growth, metabolism, and tissue repair. Several genes—including IL6, HGF, TNF, VWF, KITL, and MITF—were increased approximately two-fold from 5mo to 24mo. Elevated IL6 and TNF expression is consistent with aging-related inflammation and activation of inflammatory regulatory pathways (Bradly 2008). In contrast, NT5E and NGFR levels decreased nearly two-fold in 24mo compared to 5mo. NT5E contributes to tissue homeostasis and cell survival; thus, its reduced expression may reflect declining regenerative capacity with age (Fang et al. 2021). Similarly, downregulation of NGFR has been linked to age-related conditions such as osteoarthritis (Zhao et al. 2024).

Through RNA-seq analysis, we identified differentially expressed genes (DEGs) that provide insight into the molecular mechanisms underlying age-associated alterations in MSCs derived from 5mo, 12mo, and 24mo female C57BL/6J mice. Our results highlight several major pathways affected by aging, including metabolic dysfunction, extracellular matrix (ECM) remodeling, impaired immune signaling, cellular senescence, and dysregulated differentiation potential. The transition from 5mo to 12mo appears to represent an early phase of MSC aging, marked by relatively subtle gene expression changes. Notably, few biological processes were enriched among upregulated genes during this period, suggesting that early aging may not yet elicit significant functional disruption. However, downregulated DEGs were significantly associated with pathways involved in cell-matrix adhesion, integrin-mediated signaling, and innate immune responses. These functions are vital for MSC interactions within their niche, enabling tissue repair and immunomodulation. Integrin-mediated adhesion plays a central role in transmitting extracellular cues to intracellular pathways that govern MSC proliferation and differentiation (Hynes, 2002; Barczyk et al. 2010). Early downregulation of these genes may predispose MSCs to impaired tissue integration and reduced regenerative potential. Furthermore, the suppression of immune-related genes may reflect early immunosenescence, which contributes to the age-associated decline in immune function (Franceschi et al. 2018).

Further our RNA-seq data showed MSCs from 24mo vs 5mo mice exhibited more extensive transcriptomic remodeling. Key metabolic pathways, including glycolysis and gluconeogenesis, were significantly downregulated, indicating age-associated metabolic decline. Given that MSCs rely heavily on glycolysis for energy during proliferation and differentiation (Beegle et al. 2015; Sartorelli et al. 2025), this reduction may compromise energy homeostasis, promote senescence, and impair stem cell functionality. Simultaneously, genes involved in ECM remodeling—particularly MMP9 and MMP13, responsible for collagen degradation—were upregulated. Excessive ECM degradation may undermine bone matrix integrity, contributing to skeletal aging and disorders such as osteoporosis (Nagase et al. 2006; Verma & Hansch, 2007). Additionally, MSCs showed increased expression of genes involved in cell adhesion and negative regulation of Wnt signaling. While increased adhesion may enhance niche retention, it may also impair MSC migration. Downregulation of Wnt signaling, a key pathway in osteogenesis and MSC self-renewal, suggests a shift in lineage commitment away from osteoblastogenesis toward adipogenesis (Clevers & Nusse, 2012). Finally, between 12 and 24 months, we observed reduced expression of ribosomal genes, including those encoding components of the 30S and 60S subunits. This translational repression may impair protein synthesis and contribute to stem cell dysfunction, while concurrent upregulation of DNA damage response and cell cycle genes suggests heightened genomic stress and senescence activation (Steffen & Dillin, 2016; Campisi & d’Adda di Fagagna, 2007).

Functionally, aging altered MSC differentiation capacity, with a clear shift from osteogenesis to adipogenesis. This shift is supported by increased expression of PPARG, a master regulator of adipogenesis, and decreased expression of osteogenic markers ALP and OPN. This lineage imbalance contributes to bone loss and increased marrow adiposity observed in aging (Manolagas, 2010; Moerman et al. 2004). Reduced osteogenesis may also reflect impaired Wnt signaling and disrupted metabolic support, both of which are critical for bone formation. The reduction in Ki67 expression in aged MSCs further confirms diminished proliferative activity. Ki67 is a well-established marker of cell proliferation, and its decline is associated with impaired regeneration and reduced differentiation potential (Scholzen & Gerdes, 2000). Concurrently, increased expression of p16, a cyclin-dependent kinase inhibitor and marker of senescence, indicates that aged MSCs have entered a state of growth arrest, limiting their therapeutic potential (Sharpless & Sherr, 2015; Campisi, 2013). We also observed age-associated changes in cell stiffness. These structural impairments may limit MSCs’ ability to respond to environmental cues, further contributing to their functional decline. Interestingly, we did not find large changes in F-actin cytoskeleton. F-actin architecture around the nucleus was more variable with increasing with age suggesting that nuclei but not F-actin cytoskeleton may be disproportionally affected by aging.

Mitochondrial dysfunction is a central hallmark of aging, and aging MSCs are known to develop altered mitochondrial morphology, impaired membrane potential, and reduced bioenergetic capacity (Barilani, 2022). In our study, 12mo and 24mo MSCs showed progressive increases in mitochondrial length and volume, together with non-linear changes in mitochondrial membrane potential, indicating that structural remodeling accompanies functional adaptation during MSC aging. The pronounced elongation at 12 months is consistent with increased fusion and may serve as a compensatory response to maintain bioenergetic function under stress (Youle & van der Bliek, 2012) and upregulation of fusion and membrane-remodeling regulators at 12 months such as PLD6 and STOML2, is consistent with an adaptive fusion response to maintain mitochondrial performance (Watanabe et al. 2011; Christie et al. 2011). By 24 months, there was a shift toward a partial restoration and elevation of membrane potential likely reflecting a more maladaptive, stress-associated state. RNA-seq data further indicated that by 24 months, fusion–fission pathway changes are more balanced, with decrease in fission machinery components such as FIS1 (Serasinghe & Yoon, 2008) and increase in including UCP2 (Brand & Esteves, 2005).

mTORC1 integrates nutrient, growth factor, and stress cues to regulate mitochondrial biogenesis, dynamics, and autophagy, and chronic mTOR activation has been implicated in stem cell aging, whereas partial inhibition can enhance mitochondrial quality control (Saxton & Sabatini, 2017) The 12mo profile, characterized by higher expression of upstream PI3K and translational regulators such as AKT1 (Manning & Toker, 2017) and TSC2 (Huang & Manning, 2008) while the 24mo profile shows higher representation of negative regulators such as TSC1/2 and PTEN (Salmena et al. 2008) and reduced AKT–EIF4E axis components indicate a muted mTOR state (Magnuson et al. 2012)

Genome-maintenance related gene expression data indicates that mtDNA replication and repair pathways are also dynamically regulated with age. Upregulation at 12mo of genes involved in mtDNA processing and repair, such as RNASEH1, DNA2, SSBP1, and POLG, is consistent with an adaptive response to increasing oxidative damage, aiming to preserve mitochondrial genome integrity. Decrease of mitochondrial ribosomal components and repair factors seen in 24mo old MSCs including MRPL proteins and RRM2B, suggest a more persistent impairment of mtDNA. Oxidative phosphorylation is the main source of cellular energy, initiating majority of ATP required to support all essential cellular process. Aging progressively impairs oxidative phosphorylation through damage to mitochondrial components, impaired electron transport efficiency, reduced ATP production and increased ROS leakage (Choksi & Papaconstantinou, 2008, Xu et al. 2025). Our RNA profile data showed increased oxidative phosphorylation related genes expression at 12mo compared to 5mo, which considered to represent a compensatory response to the initiation of mitochondrial dysfunction. Previous studies have shown that acute stress can trigger an accelerated oxidative phosphorylation, which subsequently leads to increased free radical generation (Kowalczyk et al. 2021, Brace et al. 2016). On the other hand, at 24 mo compared to 5mo, most oxidative phosphorylation–related genes were downregulated. This reduction directly decreased the mitochondria’s ability to oxidize substrates and impairs the capacity for ATP synthesis. Highly downregulated genes at 24mo including COX7B2, DGUOK, MT-ND4L, NDUFAB1 responsible for reducing the ATP synthesis (Guzman et al. 2026).

The LIV experiments integrate these molecular and structural findings into a functional test of mechanosensitivity. In young (5-month) MSCs, LIV induced a strong increase in mitochondrial length and volume with a concomitant reduction in mitochondrial number, consistent with robust fusion and network remodeling as part of a healthy mechanotransduction response (Zhang et al. 2018). As mitochondria is emerging as key transducers of mechanical and metabolic cues in stem cells, where changes in morphology and dynamics help couple mechanical inputs to bioenergetic and transcriptional outputs (Branco et al. 2023), the markedly blunted morphological response to LIV in 12- and 24mo MSCs, despite identical mechanical stimulation, suggests that aged mitochondria have reduced ability to adapt mechanical stress, likely due to the altered fusion/fission, oxidative phosphorylation, mTOR, and genome-maintenance programs described above.

Finally, the non-linear behavior of mitochondrial membrane potential in response to both aging and LIV strengthens the interpretation that elevated potential in older MSCs reflects a stressed state rather than improved function. A decrease in membrane potential in 5mo and 12mo old MSCs after LIV may represent transient uncoupling and enhanced turnover, processes associated with healthy quality control, whereas the increase in membrane potential observed in 24mo MSCs at baseline and following LIV resembles hyperpolarization states linked to impaired mitophagy and ROS accumulation in aged cells (Zhang et al. 2020). Together, these findings suggest that 12mo MSCs occupy a partially compensated state with increased fusion and preserved, albeit reduced, mechanosensitivity, whereas 24mo MSCs transition to a state characterized by altered signaling, impaired genome maintenance, reduced oxidative phosphorylation process, limited structural adaptability, and a stress-like hyperpolarization response to mechanical stimulation.

Here we established a reliable *in vitro* assay system to investigate age-associated alterations in primary MSCs derived from mice bone. Our findings demonstrate that aging impairs MSC function through a combination of transcriptional reprogramming, enhanced cellular senescence, and mitochondrial dysfunction. These results underscore the multifactorial nature of MSC aging and highlight the need for targeted strategies to preserve MSC functionality and potentiality in aging tissues.

## Author Contributions

Omar Faruq: concept/design, data analysis/interpretation, manuscript writing. Nina Nikitina: concept/design, data analysis/interpretation. Scott Birks: concept/design, data analysis/interpretation. Calvin L Jones: concept/design, data analysis/interpretation, manuscript writing. Mathew Goelzer: concept/design, data analysis/interpretation, final approval of manuscript. Sean Howard: concept/design, final approval of manuscript. Anamaria Zavala: concept/design, final approval of manuscript. Nilufar Ali: concept/design, financial support, final approval of manuscript. Gunes Uzer: concept/design, data analysis/interpretation, financial support, manuscript writing, final approval of manuscript.

## Supplementary Materials

**FIGURE S1.**
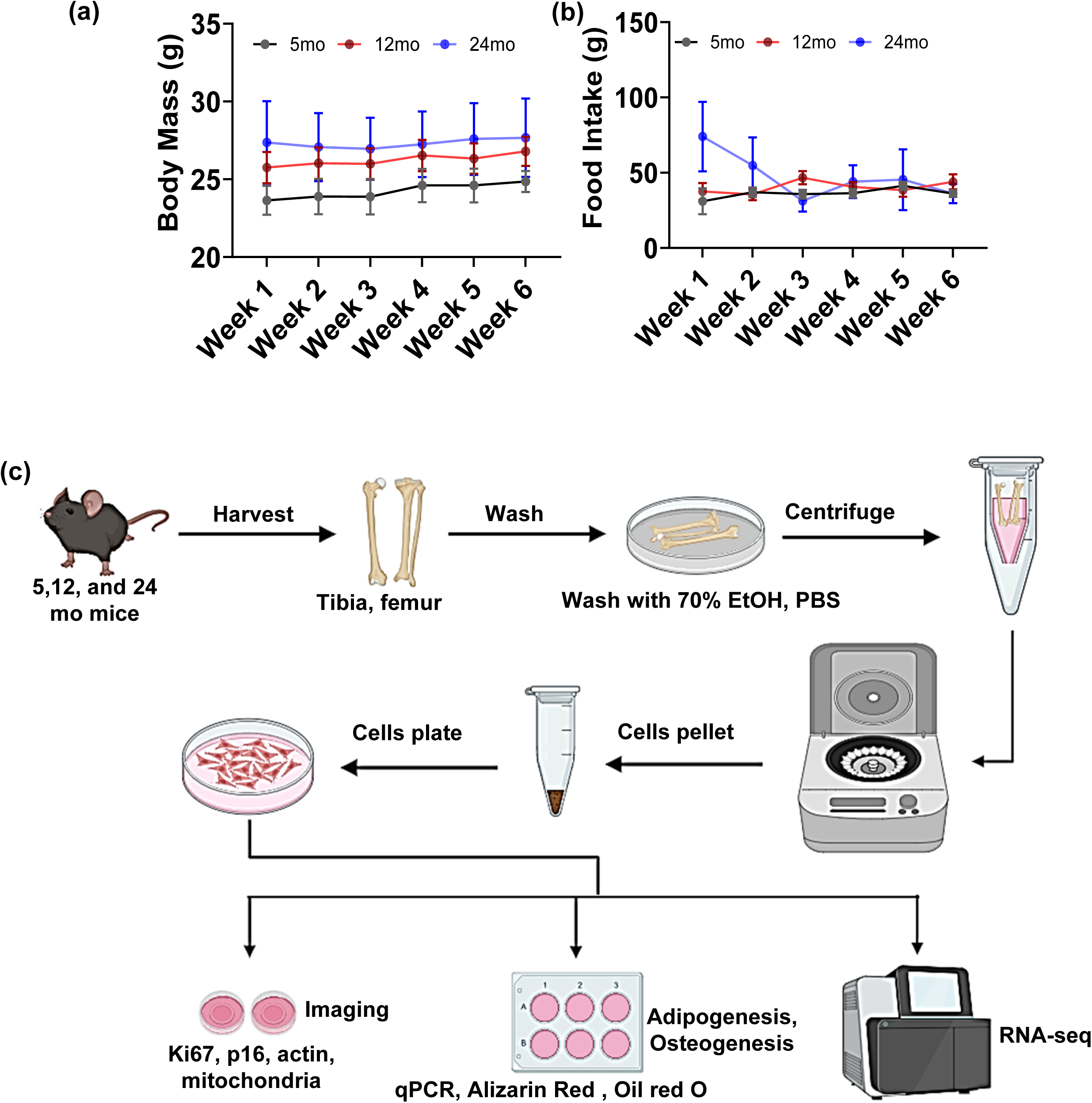
Body weight, and food intake no significant intervention in mice, workflow harvesting MSCs. (a) Body mass and (b) Food intake graph showed no significant differences among the 5mo, 12mo, and 24mo aged mice. (c) Schematic representation of workflow for isolation and characterization of aged MSCs.

**FIGURE S2.**
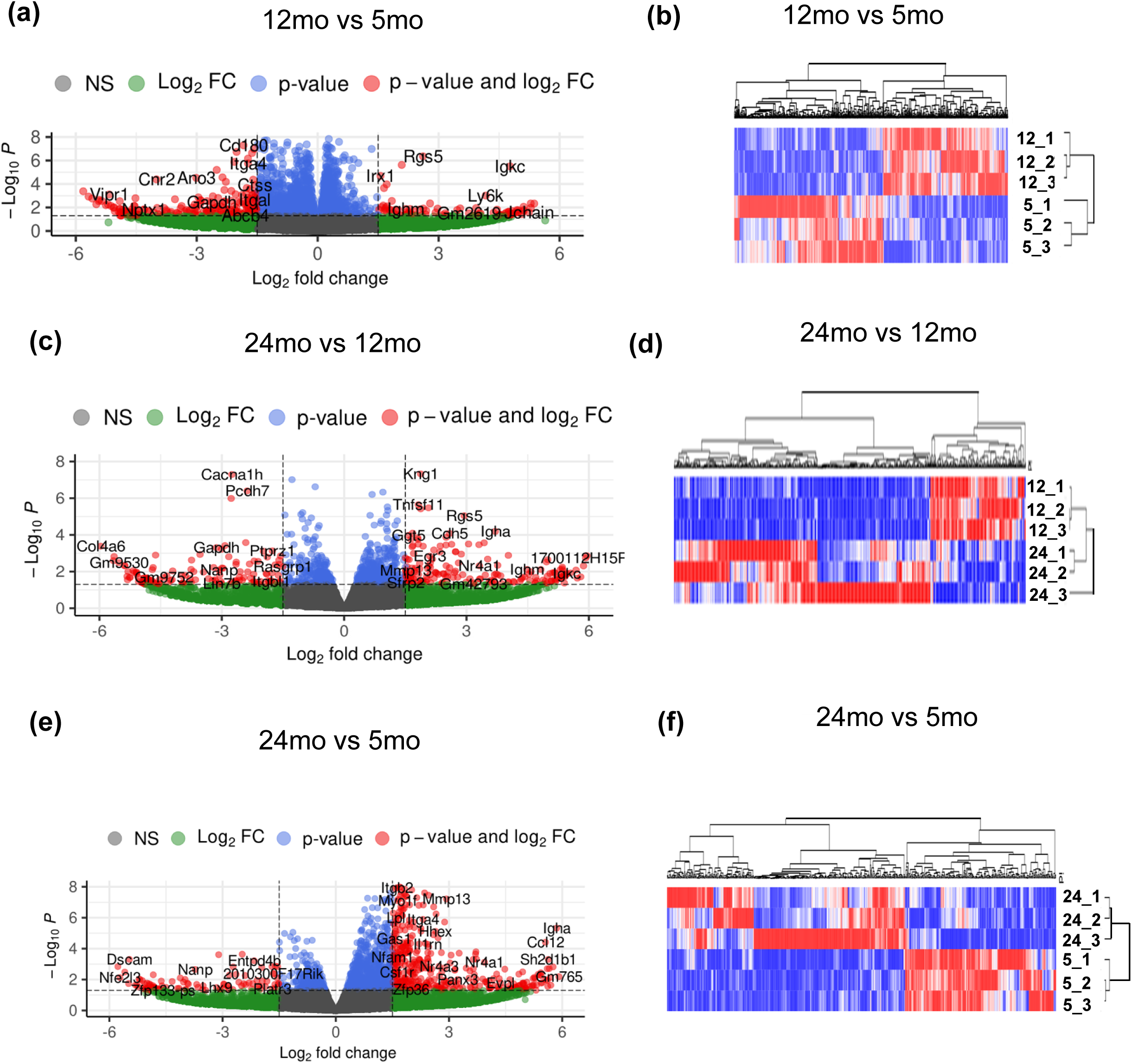
Heatmap and volcano plot compared transcriptome profiles between the different groups aged MSCs. Volcano plot compared (a) 5mo to 12mo,(c) 12mo to 24mo, and (e) 5mo to 24mo samples. Heatmap of samples after DESQ2 analysis. Hierarchical clustering compared samples and genes show transcriptome profiles separate from (b) 5mo to 12mo (d) 12mo to 24mo, and (f) 5mo to 24mo.

**FIGURE S3.**
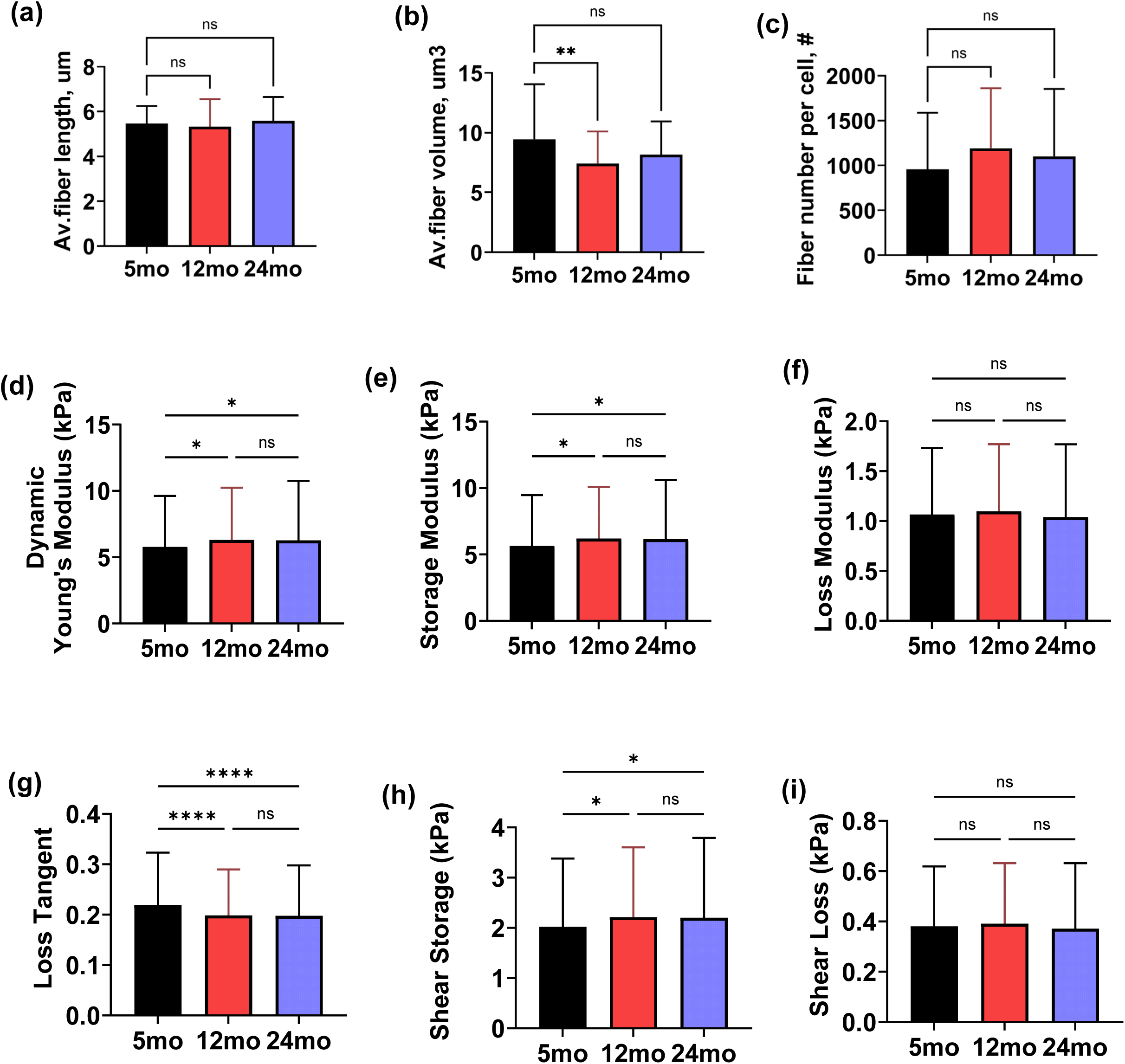
Age associated changes of actin fiber and AFM derived viscoelastic analysis of aged MSCs. Quantitative measurements include (a) avg. fiber length, (b) number and (c) volume were presented. Observed for elastic measurements, including (d) dynamic Young’s modulus (p = 0.0153 and p = 0.0301, respectively), (e) storage modulus, and (h) shear storage modulus (both p = 0.0127 and p = 0.0226, respectively). No significant differences were found in viscous measurements between age groups, including (f) loss modulus and (i) shear loss modulus. A significant decrease in (g) loss tangent was observed between 5mo and 12mo, as well as 5mo and 24mo MSCs (both p < 0.0001), with no change between 12mo and 24mo. Results are presented as mean ± SD. Significance was determined via one-way ANOVA with Tukey’s multiple comparisons. *p < 0.05, **p < 0.01, ***p < 0.001, ****p < 0.0001.

**FIGURE S4.**
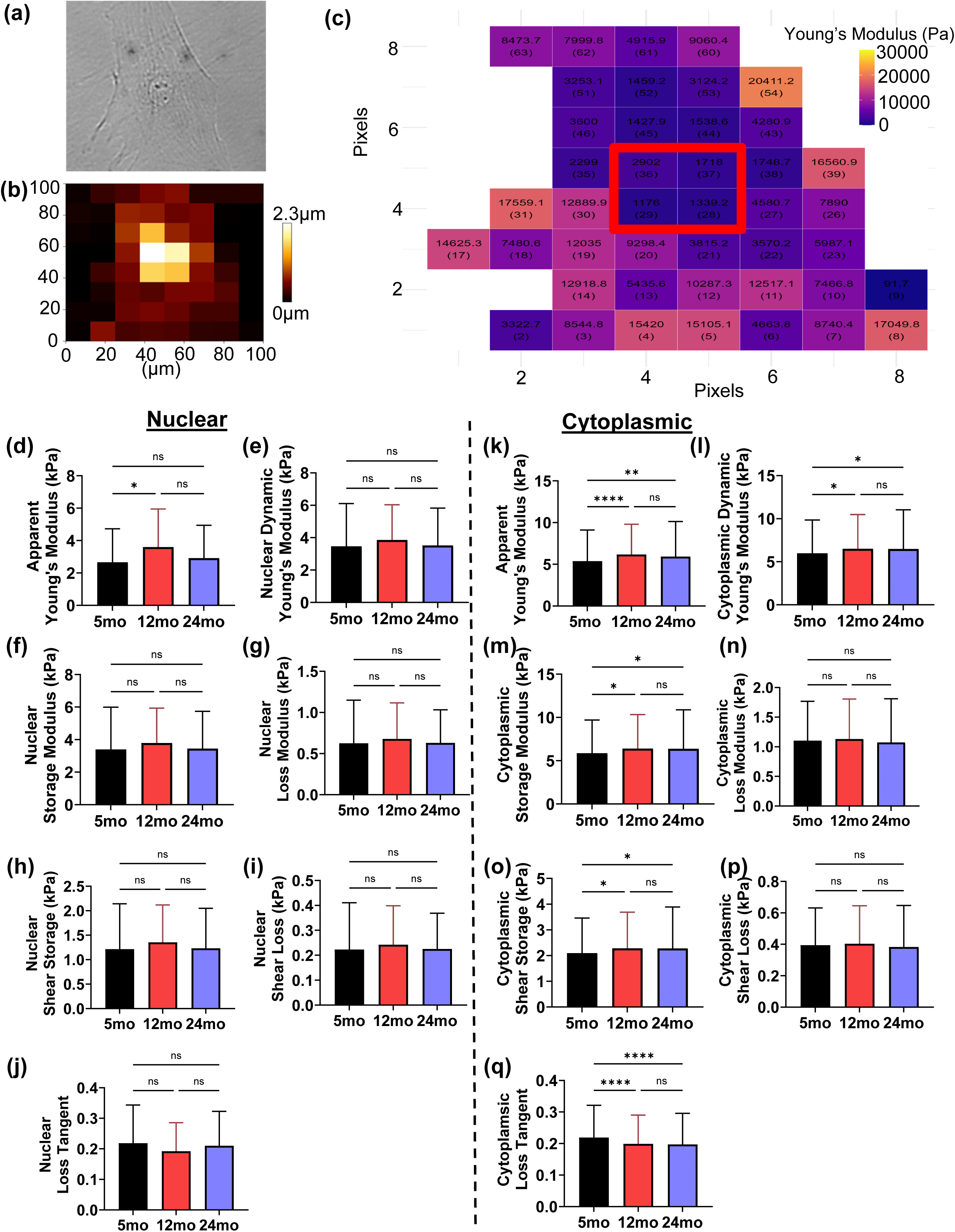
AFM derived viscoelastic analysis of aged MSCs by nuclear and cytoplasmic regions. (a) Example of brightfield image of isolated single cell with (b) height map overlay used to identify (c) nuclear region of mechanically derived force maps. (d) A significant increase in apparent Young’s modulus of the nuclear region was found between 5mo and 12mo MSCs (p = 0.0299) however no significant differences were found between 5mo and 24mo nor 12mo and 24mo. No significant differences were found in the nuclear region between any of the aged MSCs for (e) dynamic Young’s modulus, (f) storage modulus, (g) loss modulus, (h) shear storage (i) shear loss, or (j) loss tangent. Cytoplasmic region data closely mimicked whole cell data with (k) significant increases in apparent Young’s modulus observed between 5mo and 12mo, and 5mo and 24mo MSCs (p < 0.0001 and p = 0.0095, respectively), with no significant difference between 12mo and 24mo. Similar trends were observed for other elastic measurements, including (l) dynamic Young’s modulus (p = 0.0263 and p = 0.0367, respectively), (e) storage modulus, and (h) shear storage modulus (both p = 0.0221 and p = 0.0277, respectively). No significant differences were found in viscous measurements between age groups, including (f) loss modulus and (i) shear loss modulus. A significant decrease in loss tangent was observed between 5mo and 12mo, as well as 5mo and 24mo MSCs (both p < 0.0001), with no change between 12mo and 24mo. Results are presented as mean ± SD. Significance was determined via one-way ANOVA with Tukey’s multiple comparisons. *p < 0.05, **p < 0.01, ***p < 0.001, ****p < 0.0001.

**FIGURE S5.**
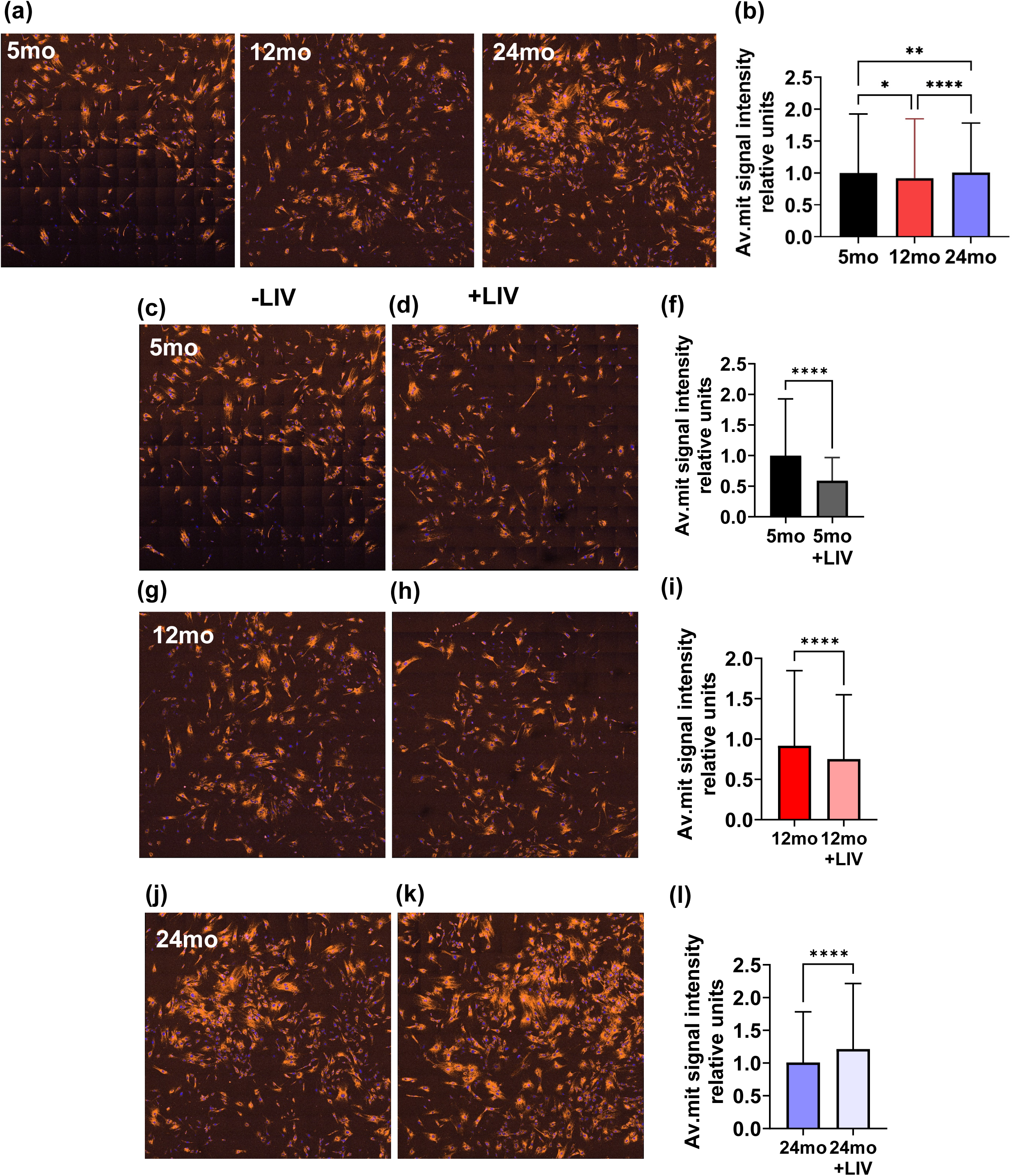
Aging increase mitochondrial membrane potential intensity and LIV modulate the intensity in each age group. (a) Images showed mitochondrial membrane potential intensity in different age group. (b) Quantitatively measurement the average mitochondrial signal intensity. Images showed the mitochondria fiber in orange for -LIV or +LIV treatment of 5mo (c, d), 12mo (g, h) and 24mo (j, k). Comparison of –LIV, +LIV treatment showed quantitively measurement of mitochondrial signal intensity for 5mo(f),12mo(i), and 24mo(l). Results represent as an average ± SD. Significance level determined via student t-test.*p<0.05, **p<0.01, ***p<0.001, ****p<0.0001

**FIGURE S6.**
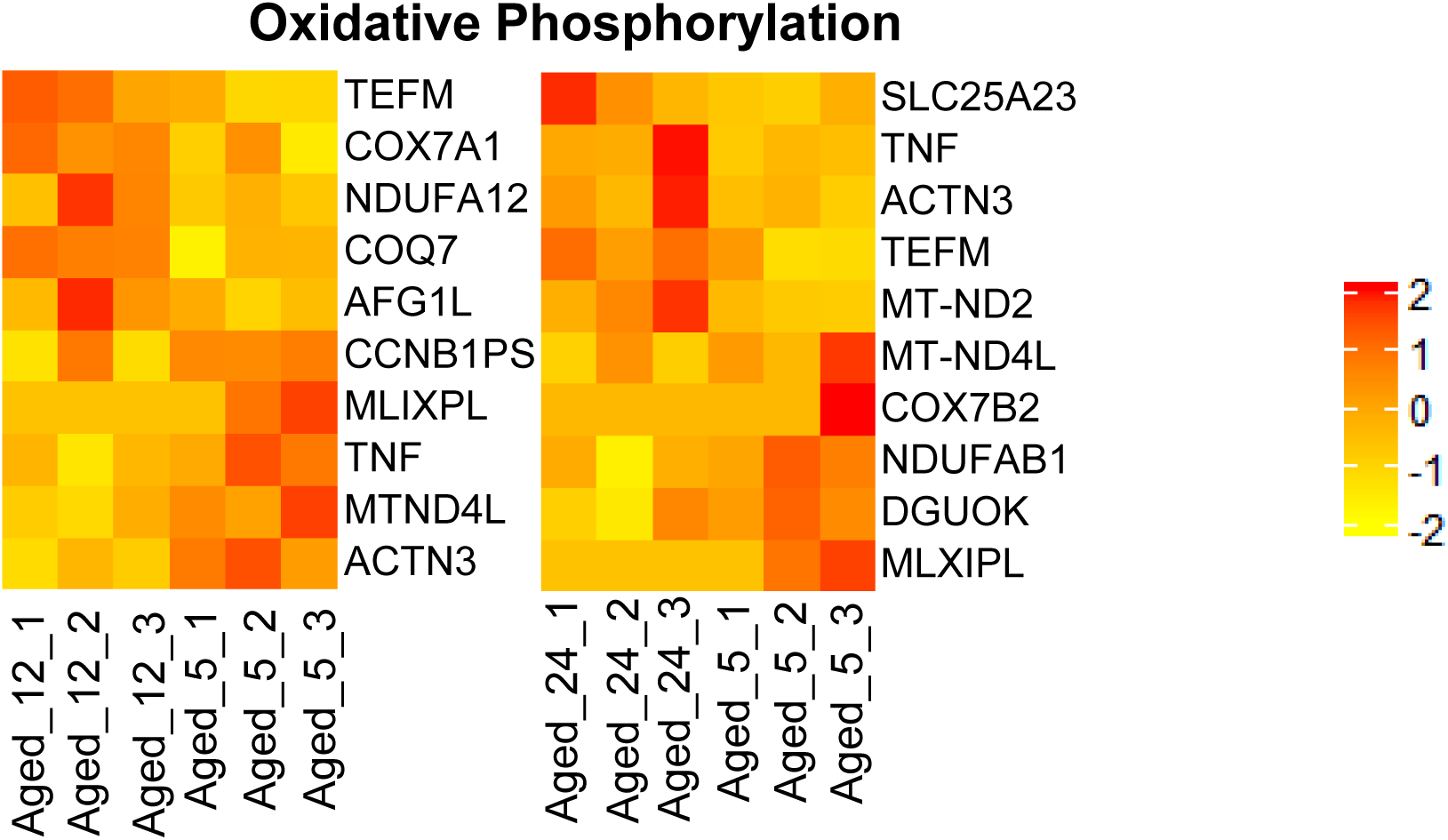
Heatmap of top 5 upregulated and downregulated Oxidative Phosphorylation related genes

**FIGURE S7.**
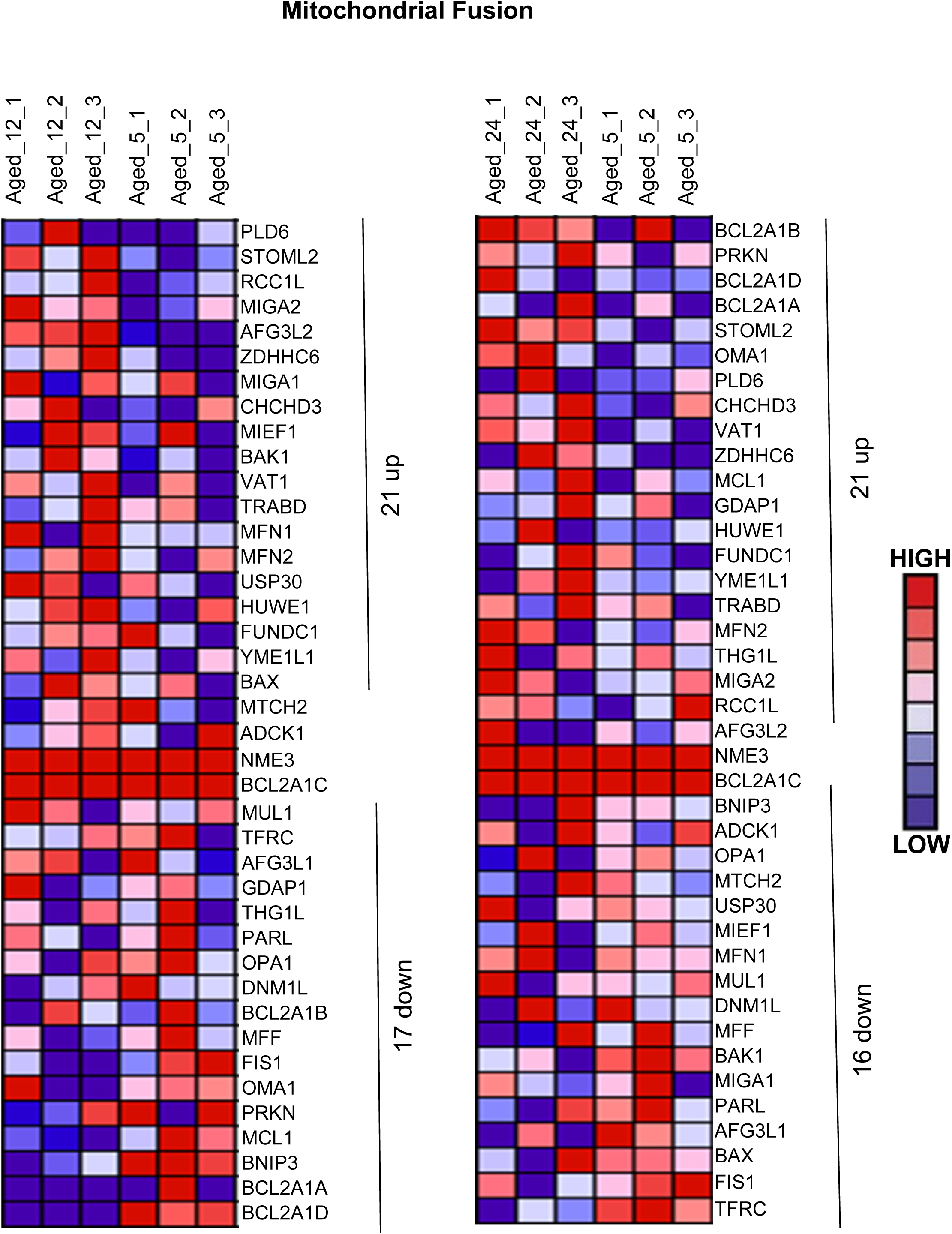
Heatmap of all upregulated and downregulated mitochondrial Fusion related genes (12mo vs 5mo; 21 upregulated and 17 downregulated, 24mo vs 5mo; 21 upregulated and 16 downregulated).

**FIGURE S8.**
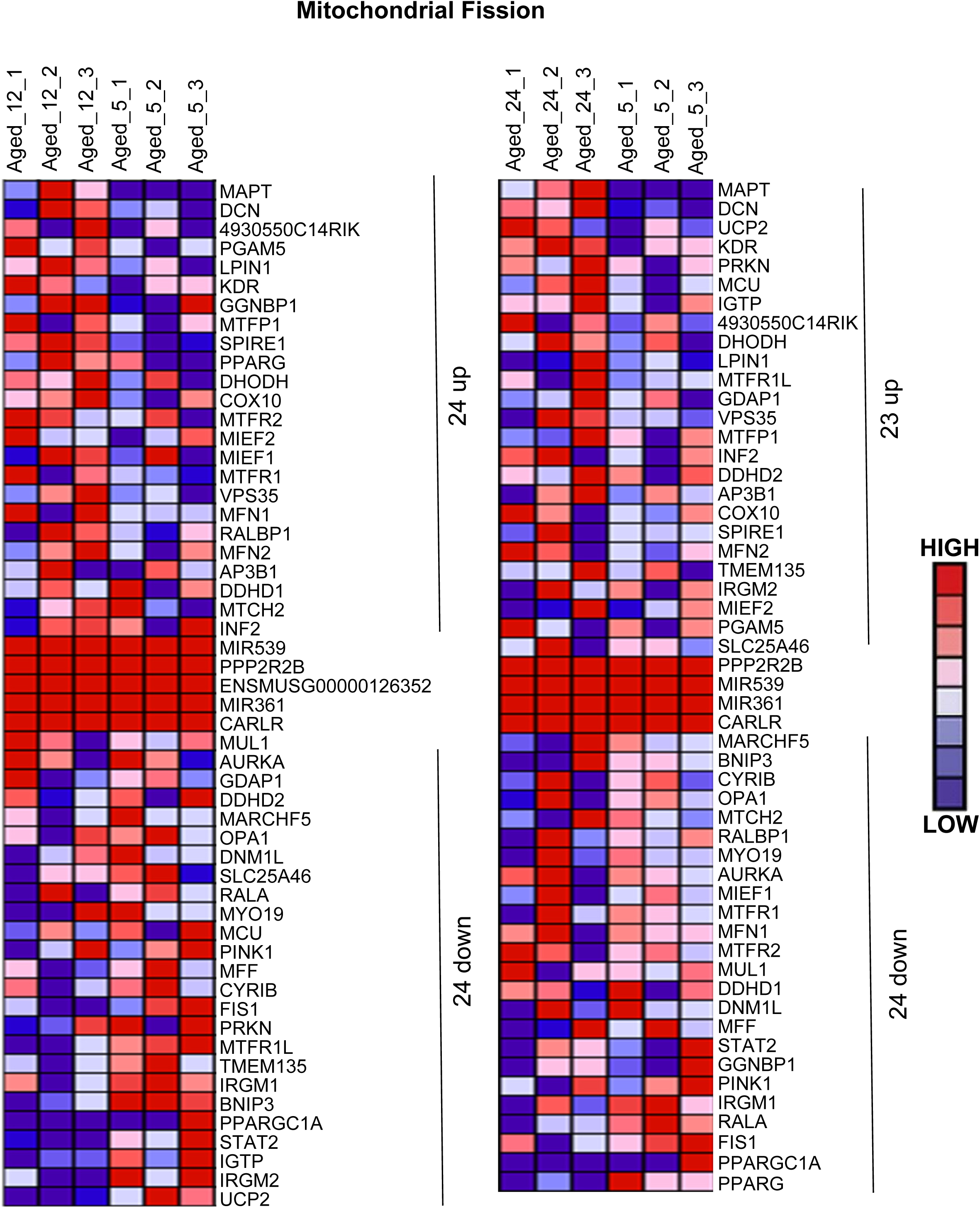
Heatmap of all upregulated and downregulated mitochondrial Fission related genes (12mo vs 5mo; 24 upregulated and 24 downregulated, 24mo vs 5mo; 23 upregulated and 24 downregulated).

**FIGURE S9.**
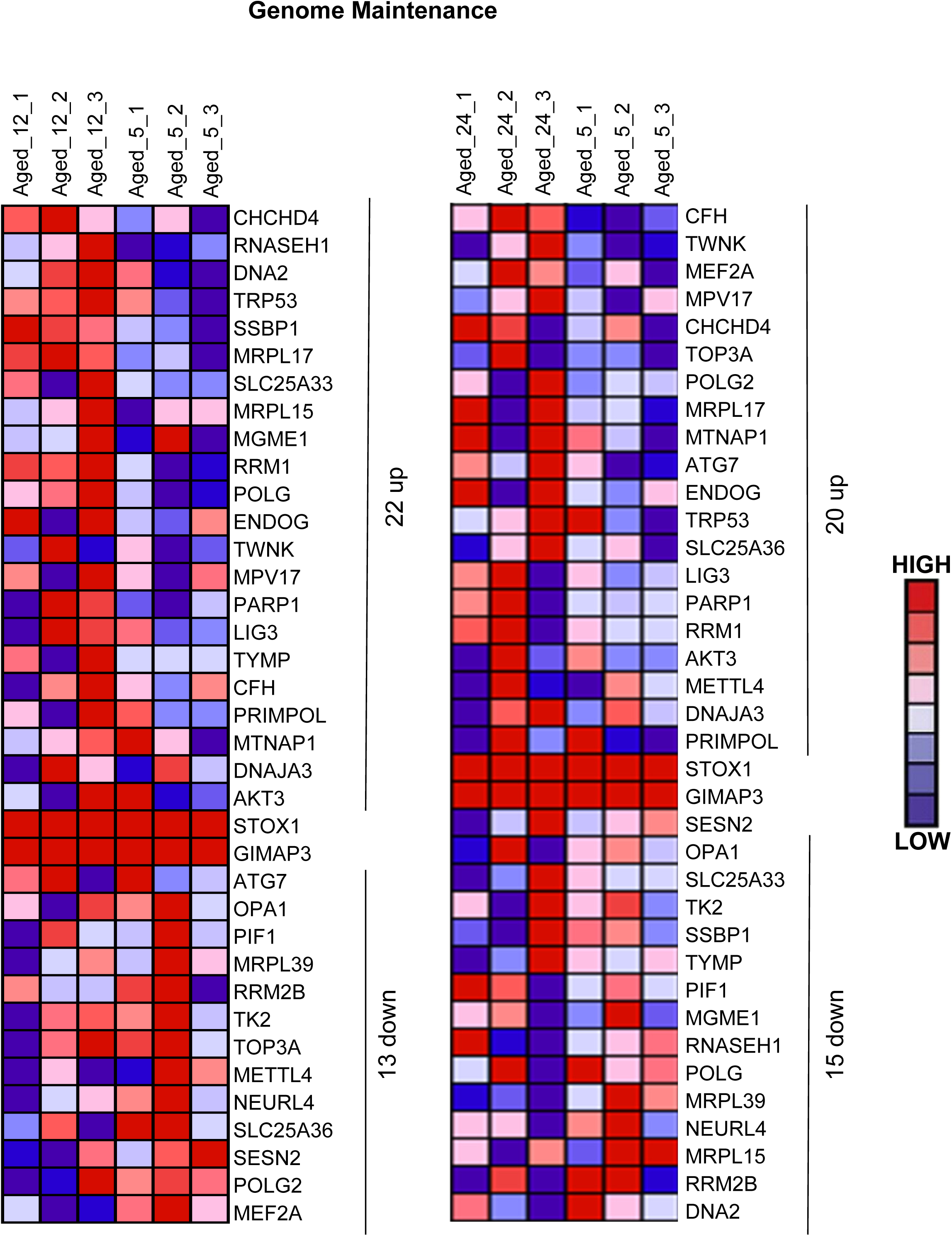
I Heatmap of all upregulated and downregulated genome maintenance related genes (12mo vs 5mo; 22 upregulated and 13 downregulated, 24mo vs 5mo; 20 upregulated and 15 downregulated).

**FIGURE S10.**
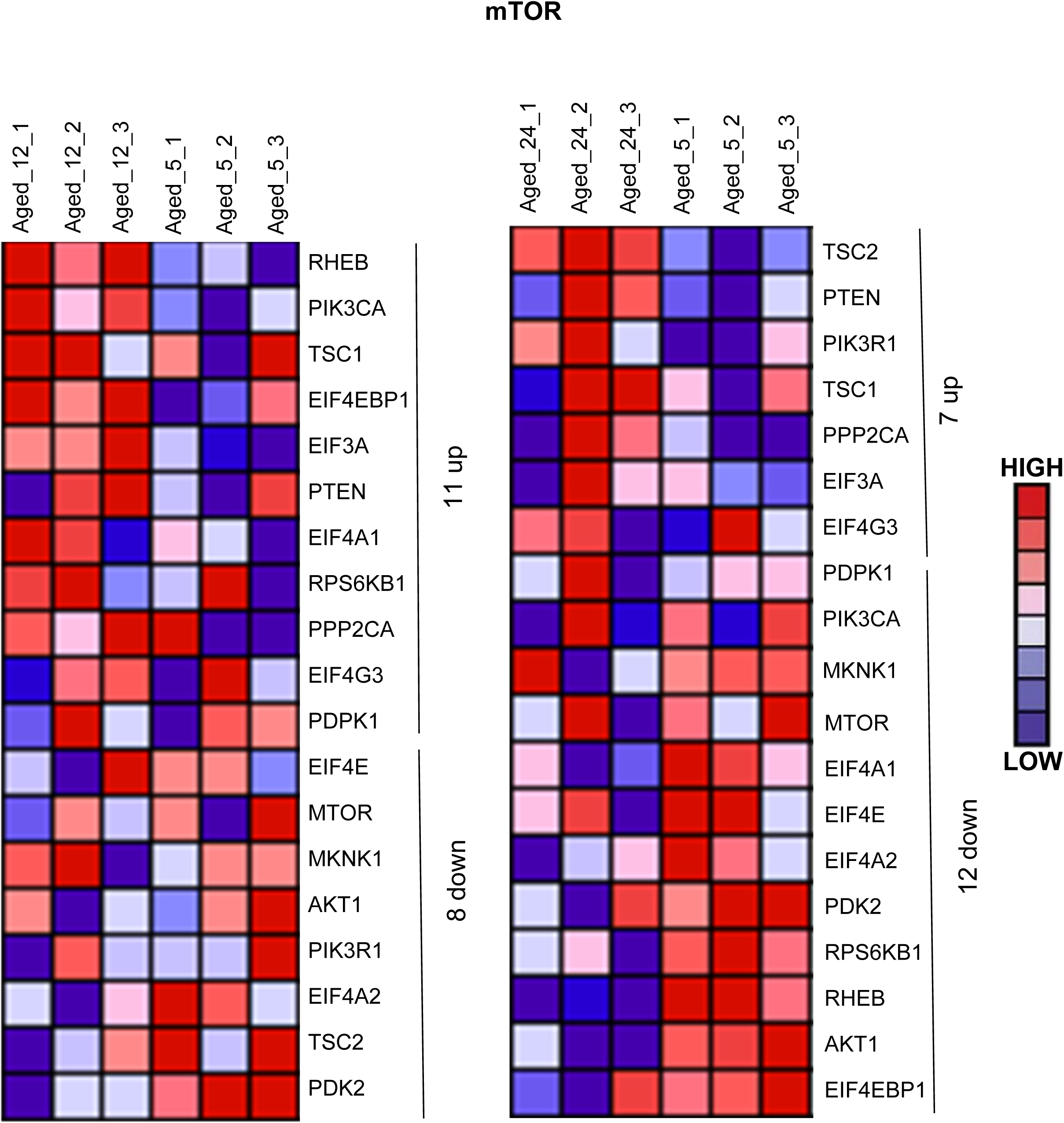
Heatmap of all upregulated and downregulated mTOR related genes (12mo vs 5mo; 11 upregulated and 8 downregulated, 24mo vs 5mo; 7 upregulated and 12 downregulated).

**FIGURE S11.**
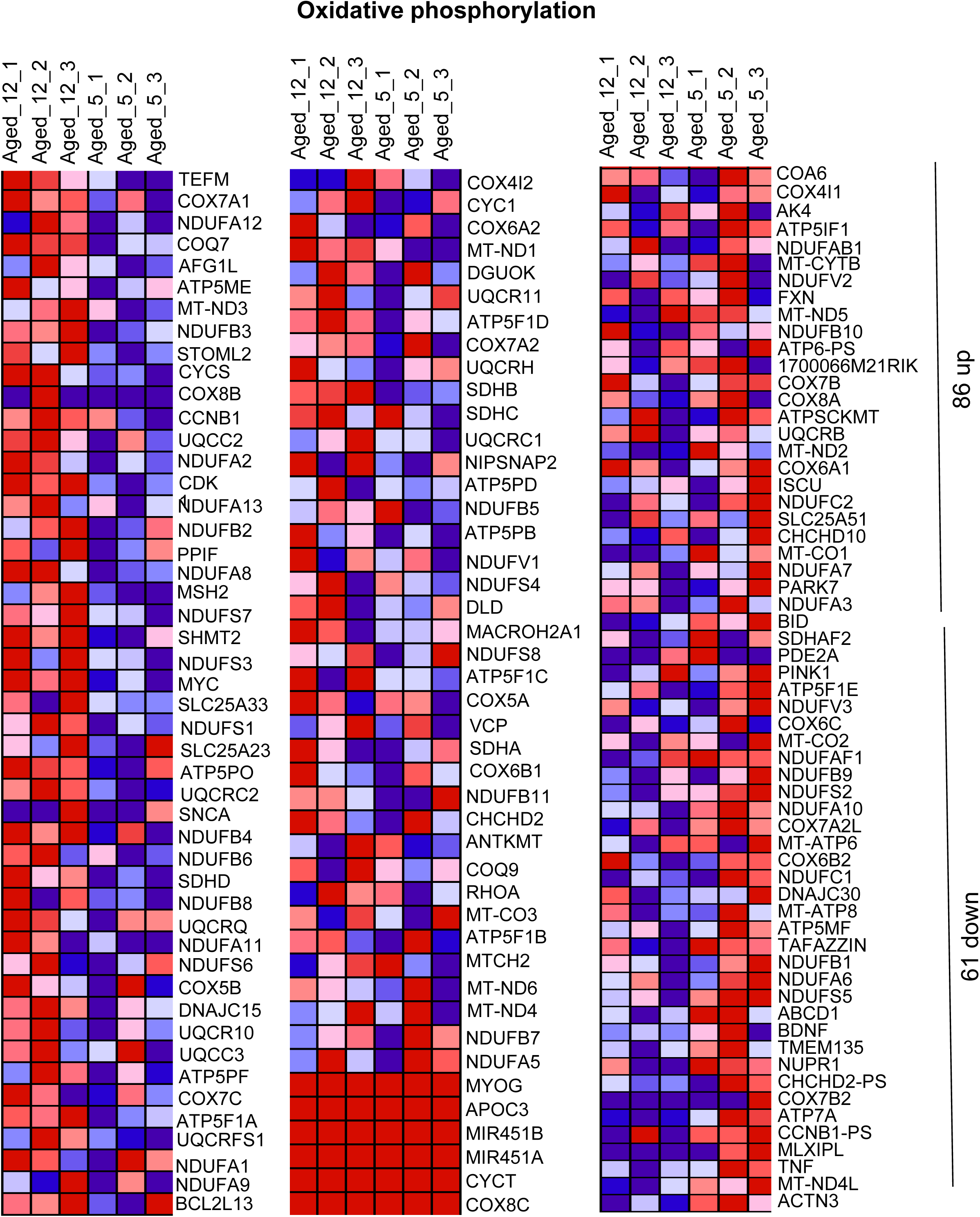
Heatmap of all upregulated and downregulated Oxidative phosphorylation related genes for 12mo vs 5mo (86 upregulated and 61 downregulated).

**FIGURE S12.**
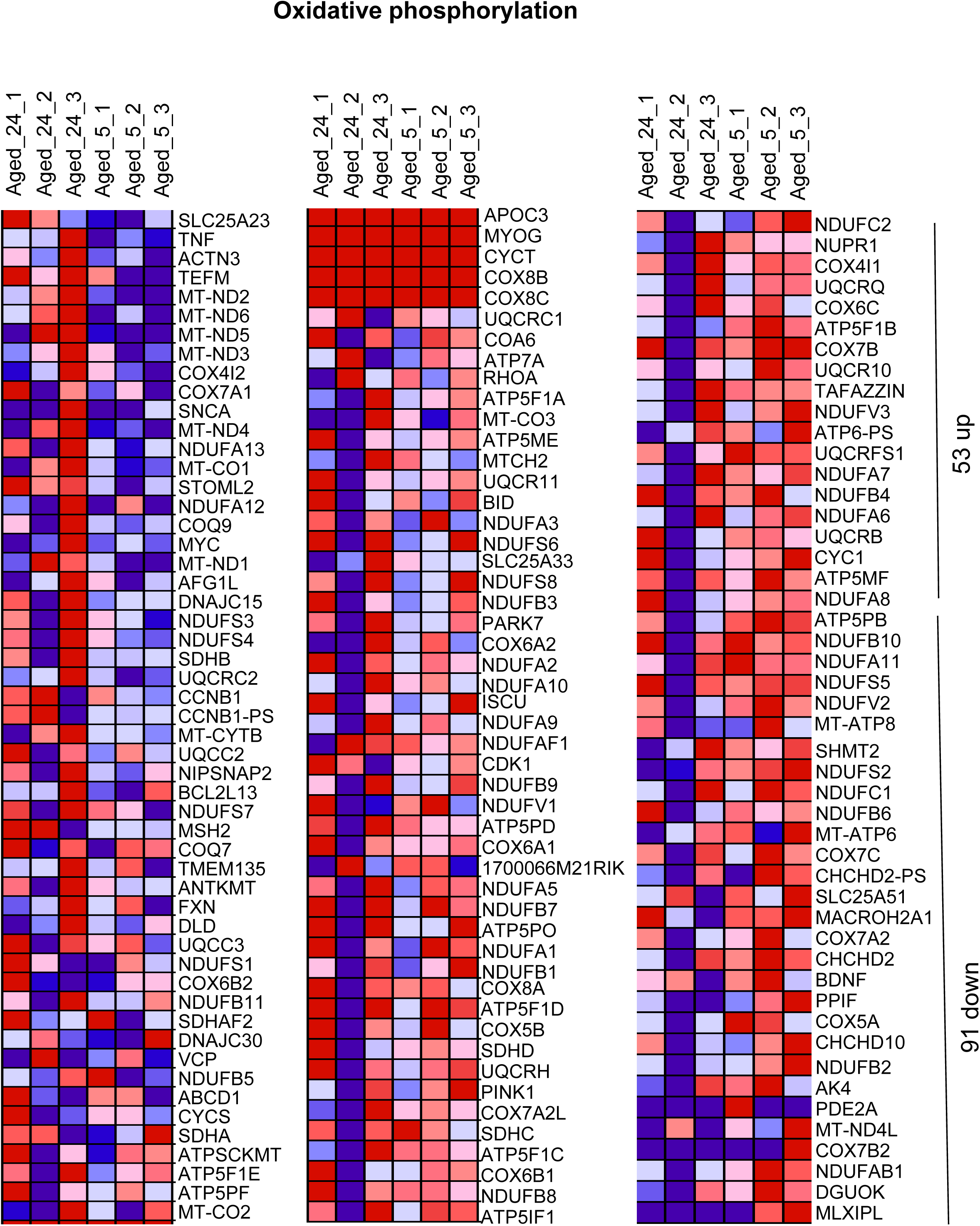
Heatmap of all upregulated and downregulated Oxidative phosphorylation related genes for 24mo vs 5mo (53 upregulated, 91 downregulated).

